# Rhomboid protease RHBDL4 promotes retrotranslocation of aggregation-prone proteins for degradation

**DOI:** 10.1101/848754

**Authors:** Josephine Bock, Nathalie Kühnle, Julia D. Knopf, Nina Landscheidt, Jin-Gu Lee, Yihong Ye, Marius K. Lemberg

**Affiliations:** Center for Molecular Biology of Heidelberg University (ZMBH), DKFZ-ZMBH Alliance, Im Neuenheimer Feld 282, 69120 Heidelberg, Germany; Center for Biochemistry, Medical Faculty, University of Cologne, Joseph-Stelzmann-Strasse 52, 50931 Cologne, Germany; Laboratory of Molecular Biology, National Institutes of Health, Bethesda, MD 20892, USA; University of Maryland School of Medicine, 670 West Baltimore Street, Baltimore, MD 21201, USA

**Keywords:** intramembrane proteolysis / rhomboid family protein / ER-associated protein degradation / prohibitin family proteins Erlin1 and Erlin2 / protein aggregates

## Abstract

Protein degradation is fundamentally important to ensure cell homeostasis. In the endoplasmic reticulum (ER), the ER-associated degradation (ERAD) pathway targets incorrectly folded and unassembled proteins into the cytoplasm for turnover by the proteasome. In contrast, lysosomal degradation serves as a failsafe mechanism for removing proteins that resist ERAD by forming aggregates. Previously, we showed that the ER-resident rhomboid protease RHBDL4, together with p97, mediates membrane protein degradation. However, whether RHBDL4 acts in concert with additional ERAD components is unclear, and its full substrate spectrum remains to be defined. Here, we show that besides membrane proteins, RHBDL4 cleaves aggregation-prone luminal ERAD substrates. Because RHBDL4 with mutations in the rhomboid domain leads to stabilization of substrates at the cytoplasmic side, we hypothesize that analogue to the homologue ERAD factor derlin, RHBDL4 is directly involved in substrate retrotranslocation. RHBDL4’s interaction with the erlin ERAD complex and reciprocal interaction of rhomboid substrates with erlins suggest that RHBDL4 and erlins form a complex that clips substrates and thereby rescues aggregation-prone peptides in the ER lumen from terminal aggregation.

## Introduction

Around one-third of all proteins enter the secretory pathway through the endoplasmic reticulum (ER), turning it into a crowded folding compartment. Even though numerous factors assist folding and complex assembly, this is an error-prone process, and misfolded polypeptides or orphan complex subunits arise that are commonly removed by the ER-associated degradation (ERAD) pathway (Christianson and Ye, 2014; Juszkiewicz and Hegde, 2018; Ruggiano et al., 2014; Wu and Rapoport, 2018). If the burden of misfolded proteins exceeds the capacity of the protein homeostasis (proteostasis) network, aggregation-prone polypeptides form clusters. Depending on the protein, these clusters consist of unstructured, amorphous aggregates or structured β-sheet amyloid fibres (Balchin et al., 2016; Breydo and Uversky, 2015). Protein aggregates cause cellular toxicity and are a hallmark of several diseases, including neurodegenerative disorders like Alzheimer’s or Parkinson’s disease (Chiti and Dobson, 2017). Clearance of large misfolded protein species in the ER is accomplished by selective autophagy (ER-phagy) or a recently described vesicular ER-to-lysosome trafficking pathway (Molinari, 2021). However, if not terminally aggregated, the best-characterized mechanism for the turnover of aberrant proteins is ERAD. Here, as part of the canonical ER quality control, misfolded proteins are recognized by a network of protein factors, including chaperones, glycan-modifying enzymes, protein disulfide isomerases, and reductases (Christianson et al., 2011).

ERAD consists of several parallel pathways that allow the removal of an exceptionally diverse set of aberrant proteins. Best understood in yeast, three major degradation routes, namely ERAD-L, ERAD-M and ERAD-C, are formed by distinct E3 ubiquitin ligase complexes that recognize proteins with lesions in the lumen, ER membrane or cytoplasm, respectively (Carvalho et al., 2006; Denic et al., 2006). Although this distinction may not be as strict in mammalian cells, defined sets of quality control factors still assist turnover of different protein classes (Bernasconi et al., 2010; Christianson et al., 2011). Glycosylated ERAD-L substrates often engage the lectins calnexin and calreticulin, α1-mannosidases (EDEM1, -2 and 3) and the disulfide reductase ERdj5 that collectively routes proteins via Sel1 to the E3 ubiquitin ligase Hrd1 (McCaffrey and Braakman, 2016; Ruggiano et al., 2014). Moreover, for turnover of soluble ERAD substrates, frequently catalytically-inactive rhomboid protease homologues referred to as pseudoproteases (Der1 and Dfm1 in yeast; Derlin1, -2 and -3 in humans) are required (Christianson et al., 2011; Greenblatt et al., 2011; Vashist and Ng, 2004). After recruitment to an E3 ubiquitin ligase complex containing a derlin protein, ERAD-L substrates are retrotranslocated across the ER membrane to reach the proteasome (Wu and Rapoport, 2018). To this end, ERAD substrates are extracted by the AAA+-ATPase p97 (Cdc48 in yeast), deglycosylated by an N-glycanase and targeted to the proteasome

(Hirsch et al., 2003; Ye et al., 2001). Work in yeast and *in vitro* suggests that for ERAD-L substrates, Hrd1 forms the core of a retrotranslocation channel (Baldridge and Rapoport, 2016; Schoebel et al., 2017). Consistent with this, a recent cryo-EM structure of the yeast Hrd1 complex revealed a sizable pore formed by two half-channels consisting of Hrd1 and Der1, which can accommodate a wide range of ERAD substrates, including bulky N-linked glycans (Wu et al., 2020). However, alternative ERAD pathways exist, as for example, degradation of activated inositol 1,4,5-triphosphate IP(3) receptors in mammals engages a Mega-Dalton (MDa) complex consisting of multiple copies of the type II membrane proteins Erlin1 and -2 and the E3 ubiquitin ligase RNF170 (Lu et al., 2011; Pearce et al., 2007; Pearce et al., 2009) but not Hrd1. As a variation to the theme, several ERAD substrates are processed by intramembrane proteases before extraction from the ER membrane (Avci and Lemberg, 2015). Accordingly, the rhomboid intramembrane protease RHBDL4 has been linked to ERAD (Fleig et al., 2012; Paschkowsky et al., 2018) that impacts key aspects of the secretory pathway such as tuning the N-linked glycosylation machinery and the rate of ER export (Knopf et al., 2020; Wunderle et al., 2016). RHBDL4 uses a bipartite substrate recognition mechanism to select certain membrane proteins with unstable TM domains.

Primarily, RHBDL4 recognizes positively charged residues within TM domains (Fleig et al., 2012; Paschkowsky et al., 2018), which destabilize the TM helix and act as a degradation signal (degron) of ERAD-M substrates (Bonifacino et al., 1990). As a second layer of control, substrate recognition occurs through a conserved ubiquitin-interacting motif at the cytosolic C-terminal tail of RHBDL4 (Fleig et al., 2012). Therefore, RHBDL4 does not solely rely on one recognition mechanism. Rather, it integrates different information, including substrate ubiquitination, before it performs the irreversible action of cleavage. What features determine whether a protein enters a classical ERAD pathway or is first cleaved by RHBDL4 or another ER protease is unknown.

By asking what influence different proteostasis factors have on the turnover of ERAD-L substrates, we discovered that in addition to its role in ERAD-M, RHBDL4 serves as a non-canonical factor in the clearance of misfolded soluble proteins in the ER lumen. This shows that the substrate spectrum of rhomboid intramembrane proteases is more diverse than initially anticipated. Moreover, we demonstrate that for clearance of luminal substrates, RHBDL4 cooperates with the erlin complex, a putative ERAD recruitment factor for aggregation-prone peptides. Since RHBDL4 ablation increases the load of insoluble versions of its substrates, we suggest that the RHBDL4-erlin complex plays an essential role in pre-aggregate clearance from the ER lumen via dislocation of substrates into the cytoplasm and proteasomal degradation.

## Results

### A targeted siRNA screen identifies RHBDL4 as an ERAD-L component

To investigate principles of ERAD pathway selection, we transfected a soluble model ERAD substrate into Hek293T cells and analyzed its steady-state level in a siRNA screen. As model substrate, we generated a truncated version of the major histocompatibility complex (MHC) class I heavy chain of 202 amino acids (MHC202), which comprises an antiparallel β-sheet and two α-helices formed by a tandem repeat of the so-called α1 and α2 domains (Figure 1A). Based on the primary sequence and crystal structure of the MHC ectodomain (Bulek et al., 2012), we predicted that the soluble MHC202 truncation forms an unstable protein containing one N-linked glycan and one disulfide bridge that exposes an extensive hydrophobic surface (Figure 1A, bottom panel right). For cell-based screening, we tested p97 that is invariant for retrotranslocation of ERAD substrates and 40 proteins that are in the ERAD protein interaction network (Christianson et al., 2011; Christianson and Ye, 2014).

**Figure 1.**
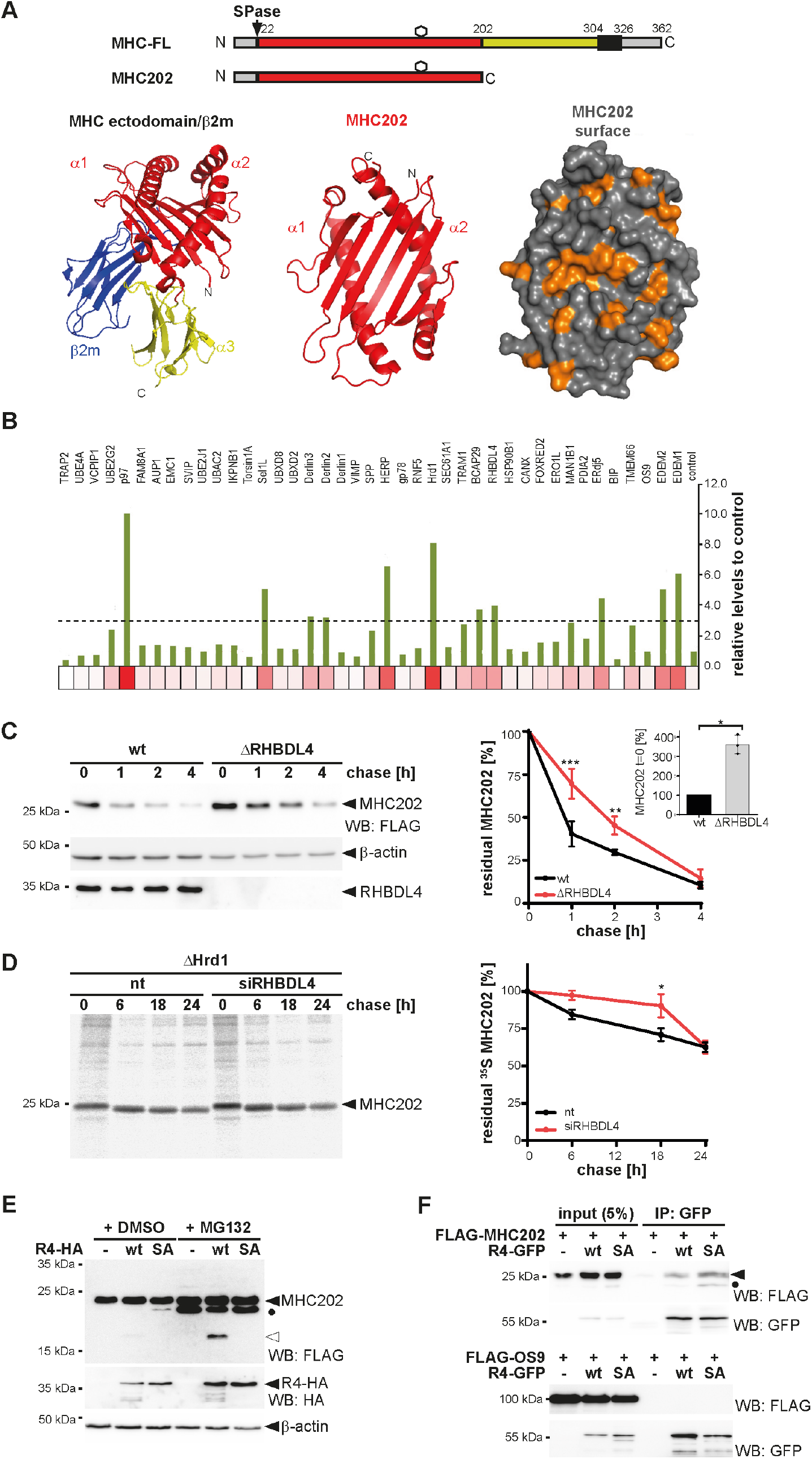
RHBDL4 contributes to the efficient turnover of a soluble ERAD substrate. **(A)** Schematic representation of full-length MHC class I heavy chain (MHC-FL) and the truncated mutant MHC202 used in this study. Black box, TM domain; hexagon, site for N-linked glycosylation; SPase, signal peptidase. The lower panel shows the crystal structure of the MHC ectodomain in complex with β2-microglobulin (β2m) taken from the atomic coordinates 3UTQ.pdb omitting the peptide ligand in the α1-α2 groove. The region comprising MHC202 is highlighted in red and shown as bottom view (middle panel). Due to the C-terminal deletion of the α3 domain, a cluster of several hydrophobic residues is exposed in MHC202 as highlighted in orange in the surface representation of the bottom view (right panel). **(B)** Targeted siRNA screen identifies non-canonical ERAD components that contribute to MHC202 turnover in addition to the Hrd1 retrotranslocation complex. Heat map of MHC202 steady-state levels using siRNA pool #1 corresponding to Figure Figure 1 – figure supplement 1A (n=1, see Supplementary file 1 for biological replicates). The threshold was set to at least three-fold change in MHC202 steady-state level. **(C)** MHC202 is stabilized in RHBDL4 knockout cells (ΔRHBDL4) when compared to wild-type Hek293T cells (wt). Turnover was evaluated 24 h post-transfection of MHC202 by adding cycloheximide (CHX) and harvesting at the indicated time points. Western blot (WB) of three independent experiments were quantified and shown in the right panel (means ± SEM, n=3; ∗p ≤ 0.05, ∗∗p ≤ 0.01; ∗∗∗p ≤ 0.001 (two-way ANOVA). Inset bar graph shows MHC202 steady state level at time point t=0 (means ± SEM, n=3; ∗p ≤ 0.05 (Student’s t-test )). β-actin was used as a loading control. **(D)** MHC202 is stabilized in Hrd1 knockout cells (ΔHrd1) for up to 24 h as analysed by metabolic label pulse-chase. RHBDL4 siRNA knockdown additionally stabilizes MHC202 significantly. The right panel shows the quantification of autoradiograms of three independent experiments (means ± SEM, n=3, ∗p ≤ 0.05 (two-way ANOVA)). **(E)** Hek293T cells were co-transfected with N-terminally FLAG-tagged MHC202 and either an empty vector (-), HA- tagged RHBDL4 (R4-HA) wt, or the catalytic inactive SA mutant. RHBDL4 generates an 18-kDa N-terminal cleavage fragment (open triangle) that is degraded by the proteasome as shown by increased steady-state level upon MG132 treatment (2 μM) compared to vehicle control (DMSO). The ectopically expressed catalytic SA mutant competes with endogenous degradation pathways and stabilizes deglycosylated full-length MHC202 (filled circle). β-actin was used as a loading control. **(F)** GFP-tagged RHBDL4 (R4-GFP) co-immunoprecipitates FLAG- tagged MHC202, but not FLAG-tagged OS9. Filled triangle, glycosylated MHC202; filled circle, deglycosylated MHC202; IP, immunoprecipitation. Data information: For clarity, for panels C-E representative experiments of 3 independent replicates are shown.

While knockdown of p97 and the E3 ligase Hrd1 showed the strongest MHC202 steady-state increase, nine other candidates also showed a strong effect (Figure 1B, Figure 1 – figure supplement 1A and Supplementary file 1). Among those were the Hrd1-associated ERAD factors Herp, Derlin2/3, Sel1, the α1-mannosidases EDEM1/2, and the disulfide reductase Erdj5. However, knockdown of the lectin OS9, which typically targets glycoprotein ERAD substrates to Sel1L did not alter MHC202 levels (Figure 1B, Figure 1 – figure supplement 1A and Supplementary file 1), indicating a redundant function of the paralogue XTP3B that has been observed for certain other glycoproteins (van der Goot et al., 2018). While all these factors are known to be involved for recognition and degradation of ERAD-L substrates (Christianson et al., 2011), we also observed that knockdown of the putative membrane-integral ER quality control factor Bap29 (Abe et al., 2009) and the rhomboid intramembrane protease RHBDL4 (Fleig et al., 2012) caused a modest increase of the MHC202 steady-state level. These factors had not been linked to Hrd1-mediated ERAD (Christianson et al., 2011), indicating that also non-canonical factors contribute to MHC202-clearance.

Intramembrane proteases are commonly believed to cleave only membrane-integral proteins, but exceptions are known (Kühnle et al., 2019). We, therefore, set out to characterize the unexpected role of RHBDL4 in MHC202 turnover. First, we confirmed that knockdown with two independent targeting sequences elevated MHC202 steady-state levels (Figure 1 – figure supplement 1B). The role of RHBDL4 in MHC202 turnover was further confirmed by cycloheximide chase in RHBDL4 knockout Hek293T cells, in which the half-live of MHC202 turnover increased from less than one hour to approximately two hours (Figure 1C).

However, inhibition was only partial, indicating that a redundant ERAD pathway targets MHC202 for degradation. Hence, we presume that induction of the ER unfolded protein response (UPR) observed upon RHBDL4 ablation (Fleig et al., 2012) masks the RHBDL4 knockout phenotype to a certain extent by upregulating alternative degradation routes. We generated Hrd1 knockout cells to investigate further whether RHBDL4 might act in parallel to the canonical Hrd1 pathway. The Hrd1 substrate null Hong Kong mutant of α1-antitrypsin (NHK) (Christianson et al., 2011) was fully stabilized in Hrd1 knockout cells, but the phenotype was reverted upon reexpression of Hrd1 (Figure 1 – figure supplement 1D). We then used metabolic pulse label chase analysis to follow up on the MHC202 degradation kinetics in Hrd1 knockout cells. While for newly synthesized MHC202 in Hek293T wild-type (wt) cells an initial fast decay was observed, a fraction persisted for a longer time different to the cycloheximide chase, increasing the apparent half-life to approximately 2 h (Figure 1 – figure supplement 1C). In contrast, in Hrd1 knockout cells, most MHC202 (∼65%) was stable even up to 24 h (Figure 1D), indicating that Hrd1 is the prime degradation route for MHC202. Despite that prominent stabilization of MHC202 in Hrd1 knockout cells, the 24 h chase revealed that approximately 35 % of MHC202 was still degraded within 24 h (Figure 1D).

However, knockdown of RHBDL4 in this genetic background leads to a further stabilization of MHC202 for up to 18 h (Figure 1D), indicating that RHBDL4 acts as a second slower degradation route. Taken together, these additive effects show that two independent pathways can remove MHC202 with different kinetics, namely canonical Hrd1-dependent retrotranslocation, and a so-far unrecognized pathway that relies on RHBDL4. The mechanism that removes MHC202 when Hrd1 and RHBDL4 are both blocked and whether ER-phagy can compensate awaits further characterization.

To investigate whether RHBDL4 directly processes MHC202, we performed a cell-based rhomboid gain-of-function cleavage assay (Fleig et al., 2012). Consistent with such a direct role of rhomboid-catalyzed cleavage in MHC202 clearance, overexpression of RHBDL4 (wt) but not its catalytic inactive serine-144-alanine mutant (RHBDL4-SA) generated an N-terminal fragment with an apparent molecular weight of 18 kDa (Figure 1E). While only traces of this cleavage product were observed in vehicle-treated cells, inhibition of the proteasome with MG132 increased its steady-state level. This result indicates that RHBDL4 generates an MHC202-cleavage fragment, which is dislocated into the cytoplasm for proteasomal degradation in an analogous manner as previously described for membrane-integral substrates (Fleig et al., 2012). Likewise, proteasome inhibition also stabilized deglycosylated full-length MHC202 (Figure 1E and Figure 1 – figure supplement 1E). Again, this shows that MHC202 is degraded by the canonical Hrd1 retrotranslocation route and an RHBDL4-dependent substrate clipping mechanism. Consequently, the full extent of RHBDL4 activity can only be seen when the downstream clearance pathway for fragments is blocked. Interestingly, overexpression of the catalytic inactive SA mutant stabilized a deglycosylated form of MHC202 even in the absence of MG132. This observation suggests that in a dominant-negative manner, the RHBDL4-SA mutant traps a partially retrotranslocated form of MHC202, exposing the glycosylation site to the cytoplasmic N-glycanase, while MHC202 is still bound to the rhomboid active site. A similar trapping effect was previously observed by overexpression of a mutant form of the rhomboid pseudoprotease Derlin1 (Greenblatt et al., 2011). Consistent with this model, the RHBDL4-SA-induced deglycosylated MHC202 species is not observed upon siRNA knockdown of N-glycanase (Figure 1 – figure supplement 1F).

Furthermore, in a co-immunoprecipitation assay from Triton X-100-solubilized cells, RHBDL4-SA co-purified the glycosylated as well as the deglycosylated MHC202, whereas ectopically expressed FLAG-tagged OS9 was not bound (Figure 1F).

### RHBDL4-catalyzed cleavage of MHC202 and p97-mediated extraction are coupled

The observation that RHBDL4-SA functionally interacts with deglycosylated MHC202 indicates that rhomboid-catalyzed cleavage and protein dislocation into the cytoplasm are linked. As the ER-integral metalloprotease ZMPSTE24 (Ste24 in yeast) has been shown to clear polypeptide chains that got stuck in the Sec61 translocon channel during post- translational translocation (Ast et al., 2016), we decided to analyse the localization of MHC202 relative to the ER lumen before cleavage. As shown above, Endo H analysis reveals that the RHBDL4-generated N-terminal MHC202 fragment is glycosylated (Figure 1 – figure supplement 1E), indicating that it is formed in the ER lumen. To also discriminate between a putative translocation intermediate with the C-terminus facing the cytoplasm and a fully translocated protein, we generated an MHC202 construct harbouring an additional glycosylation site (K197N) in the C-terminal region (Figure 2 – figure supplement 1A). We reasoned that only fully translocated MHC202 would be glycosylated at this site. Western blot analysis of MHC202-K197N co-expressed with RHBDL4 showed an Endo H-sensitive C- terminal fragment (Figure 2 – figure supplement 1A). Consistent results were obtained with an MHC202 mutant with a single C-terminal glycosylation site only (Figure 2 – figure supplement 1B), corroborating that RHBDL4 cleaves fully translocated MHC202. Consistent with this, RHBDL4 did not cleave an artificially designed ERdj3-GFP-chimera that cloggs the Sec61 translocon (Figure 2 – figure supplement 1C) and which previously had been shown to be cleaved by ZMPSTE24 (Ast et al., 2016). To further prove that RHBDL4 deals with ERAD-L substrates, we performed a protease protection assay of isolated microsomes.

**Figure 2.**
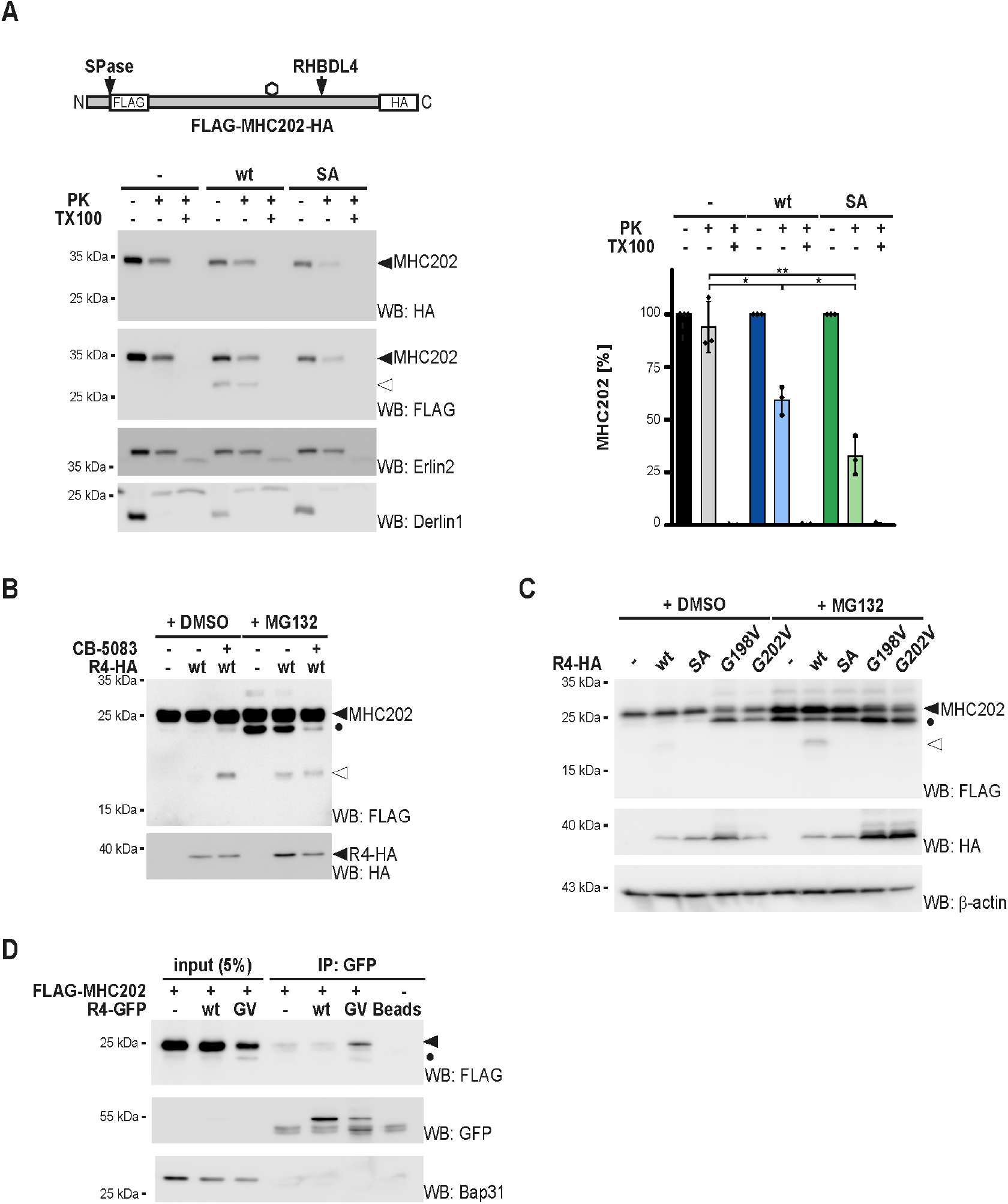
RHBDL4 cleaves MHC202 retrotranslocation intermediate and p97 facilitates dislocation into the cytoplasm. **(A)** Accessibility of MHC202 to exogenous proteinase K (PK) was analysed in ER-derived microsomes Hek293T cells were co-transfected with double-tagged HA-MHC202-FLAG and either an empty vector (-), RHBDL4 wild type (wt), or the catalytic inactive SA mutant. Hek293T-derived microsomes were incubated with PK and 1 % TritonX-100 (TX100) as indicated. R4-induced MHC202 cleavage fragment (open triangle) was protected from exogenous PK, whereas full-length MHC202 was partially accessible. Erlin2 (epitope in ER lumen) and Derlin1 (epitope in cytosol) were used as controls. Western blot (WB) HA signals of three independent experiments were quantified and are shown in the right panel (means ± SEM, n=3, ∗p ≤ 0.05; ∗∗p ≤ 0.01 (Student’s t-test). **(B)** Clearance of RHBDL4 generated cleavage product depends on p97, as shown by p97 inhibitor CB-5083 (2.5 μM) induced stabilization of the N-terminal MHC202 fragment (open triangle) even in the absence of proteasome inhibitor MG132 (2 μM). Likewise, CB-5083 reduced the appearance of the deglycosylated unprocessed form of MHC202 (filled circle) observed upon MG132 treatment, confirming that the Hrd1-dependent dislocation pathway depends on p97. Filled triangle, glycosylated MHC202; open triangle, 18-kDa N-terminal cleavage fragment. **(C)** C-terminal FLAG-tagged MHC202 is cleaved by HA-tagged RHBDL4 wt, but not by RHBDL4-G198V and RHBDL4-G202V mutants or the catalytic inactive SA mutant. β-actin was used as a loading control. Filled triangle, glycosylated full-lenght MHC202; filled circle, deglycosylated full length MHC202; open triangle, N-terminal cleavage fragment of MHC202. **(D)** GFP-tagged RHBDL4-G202V (GV) co-immunoprecipitates C-terminal FLAG-tagged MHC202. Filled triangle, glycosylated full-lenght MHC202; filled circle, deglycosylated full length MHC202; IP, immunoprecipitation. Bap31 was used as a negative control. Data information: For clarity, for panels A-D representative experiments of 3 independent replicates are shown.

While a certain fraction of MHC202 was accessible to the protease, the RHBDL4-generated cleavage fragment was protected, confirming that it is generated in the ER lumen (Figure 2A). Consistent with this, we observed ER localization of MHC202 under RHBDL4 knockdown conditions by immunofluorescence microscopy (Figure 2 – figure supplement 1D and E). Interestingly, co-expression of the RHBDL4 SA trapping mutant significantly increased the pool of protease accessible MHC202 (Figure 2A), supporting our model of a stalled retrotranslocation intermediate.

To reach the proteasome, RHBDL4-generated cleavage fragments have to be dislocated into the cytoplasm. For this purpose, RHBDL4 recruits p97 to the ER membrane via a conserved motif (termed VBM) at its cytoplasmic C-terminus (Fleig et al., 2012; Lim et al., 2016).

Blocking this interplay leads to the accumulation of glycosylated RHBDL4-generated cleavage fragments in the ER fraction (Fleig et al., 2012). Consistent with this, the p97 inhibitor CB-5083 (Anderson et al., 2015) stabilized the 18-kDa N-terminal MHC202 fragment (Figure 2B). The addition of MG132 did not further increase recovery of the cleavage fragment, indicating that solely blocking p97 and consequent retention in the ER prevents RHBDL4 generated fragments from proteasomal clearance. Additionally, we replaced a conserved arginine in the VBM, which is required for the interaction with p97, with an alanine (R308A) (Lim et al., 2016). As shown for the p97 inhibitor treatment, co-expression of the RHBDL4-R308A mutant or an RHBDL4 deletion mutant lacking the entire binding motif (RHBDL4ΔVBM) together with MHC202 results in the stabilization of the 18-kDa N-terminal MHC202 fragment (Figure 2 – figure supplement 1F). Together with the trapping of deglycosylated MHC202 with the RHBDL4-SA active site mutant, these results indicate that RHBDL4 interacts with MHC202 during retrotranslocation, thereby generating cleavage fragments that are released into the cytoplasm where they become degraded by the proteasome. Interestingly, we observed that deletion of the entire cytoplasmic domain of RHBDL4 harbouring a conserved ubiquitin-interacting motif does not prevent RHBDL4- mediated cleavage but also stabilizes deglycosylated MHC202 (Figure 2 – figure supplement 1G). As combining this effect with the substrate trapping active site SA mutant leads to an additive stabilization of deglycosylated MHC202, we speculate that RHBDL4 also interacts with a certain fraction of its substrates in a non-proteolytic manner, as has recently been described for the bacterial rhomboid protease YqgP (Began et al., 2020). This shows a striking similarity to the derlin-mediated retrotranslocation along the Hrd1 pathway (Wu and Rapoport, 2018).

Although the exact pseudoprotease mechanism in ERAD remains to be determined, previous work showed that mutation of a strictly conserved di-glycine motif (GxxxG) in TM helix 6 of the rhomboid fold leads to a dominant-negative substrate-trapping mutant of Derlin1 (Greenblatt et al., 2011). Hence, we asked whether also the rhomboid-fold of RHBDL4 directly contributes to the retrotranslocation of MHC202 and its cleavage fragments.

Consistent with a mechanistic parallel to derlin-mediated retrotranslocation, co-expression of MHC202 with RHBDL4 mutants of the di-glycine motif, namely glycine-198-valine (G198V) or glycine-202-valine (G202V), induced deglycosylated MHC202 in the absence of MG132 (Figure 2C). In agreement with interaction, immunoprecipitation of RHBDL4-G202V co-purified a substantial amount of MHC202, both in its glycosylated and deglycosylated form (Figure 2D). For both mutants, no MHC202 cleavage fragments were observed under proteasome inhibition (Figure 2C), indicating that also for RHBDL4, the GxxxG motif is critical for its activity. This had been observed for bacterial rhomboids before (Baker and Urban, 2012). Consistent with what has been observed for the SA trapping mutant, co-expressing RHBDL4-G202V with MHC202 significantly increased its accessibility in a protease protection assay (Figure 2 – figure supplement 1H). Of note, mutating the GxxxG motif did not affect the interaction of RHBDL4 with p97 and its additional binding partners (Figure 2 – figure supplement 1I and see below), indicating that the protease forms its physiological complexes while in a dominant-negative manner stalling the retrotranslocation intermediates. Overall, these results reveal that RHBDL4 plays a previously unanticipated direct role in inducing retrotranslocation of the ERAD-L substrate MHC202 and its cleavage fragments.

The exact molecular mechanism of how the rhomboid-fold of RHBDL4 contributes to retrotranslocation remains to be investigated.

### RHBDL4 cleaves selected soluble ERAD-L substrates

Next, we asked whether also other soluble ERAD-L substrates are processed in the cell-based RHBDL4 cleavage assay. However, neither NHK (Hosokawa et al., 2003) nor an ER-retained mutant of prolactin (Prl-KDEL) (Fleig et al., 2012) were processed by ectopically expressed RHBDL4 (Figure 3A). This suggests that RHBDL4 shows substrate specificity. As a follow-up, we tested two additional ERAD substrates resembling truncated type I membrane proteins, namely RI332, a deletion of ribophorin 1 (RPN1) (Tsao et al., 1992), and a loss-of-function splice variant of the β-secretase (BACE476Δ) (Tanahashi and Tabira, 2007). BACE476Δ was cleaved by ectopically expressed RHBDL4 leading to a 50-kDa fragment that appears between the glycosylated full-length 54-kDa form of BACE476Δ and the MG132-stabilized 45-kDa deglycosylated species (Figure 3B and Figure 3 – figure supplement 1A). Interestingly, ectopic expression of RHBDL4 diminished the steady-state level of BACE476Δ and completely depleted the MG132-sensitive deglycosylated full-length 45-kDa species. This suggests that upon overexpression, RHBDL4 interacts with its substrates before they approach Hrd1 and thereby outcompetes the retrotranslocation of unprocessed BACE476Δ. Consistent with a scenario of dislocating shorter, RHBDL4- generated BACE cleavage fragments into the cytoplasm, an overexposed western blot reveals a 40-kDa BACE-peptide in response to MG132 treatment (Figure 3B). Although we previously observed that degradation kinetics in Hek293T cells were unaffected by RHBDL4 knockdown for RI332 (Fleig et al., 2012), processing of RI332 by an unknown ER protease had been observed before (Mueller et al., 2006). Consistently, co-expression of RI332 with RHBDL4 generated several RI332 fragments in the range of 25 to 35 kDa, whereas the SA mutant stabilized traces of deglycosylated unprocessed species as previously observed (Figure 3C and Figure 3 – figure supplement 1B). Remarkably, the type I membrane protein RPN1 is a native RHBDL4 substrate (Knopf et al., 2020). In addition to canonical cleavage in the TM region, RPN1 is cleaved at the same position as the truncated RI332 ERAD substrate (Figure 3 – figure supplement 1B). This indicates that substrate selection of soluble substrates occurs in a related manner to cleavage of membrane-anchored ectodomains.

**Figure 3.**
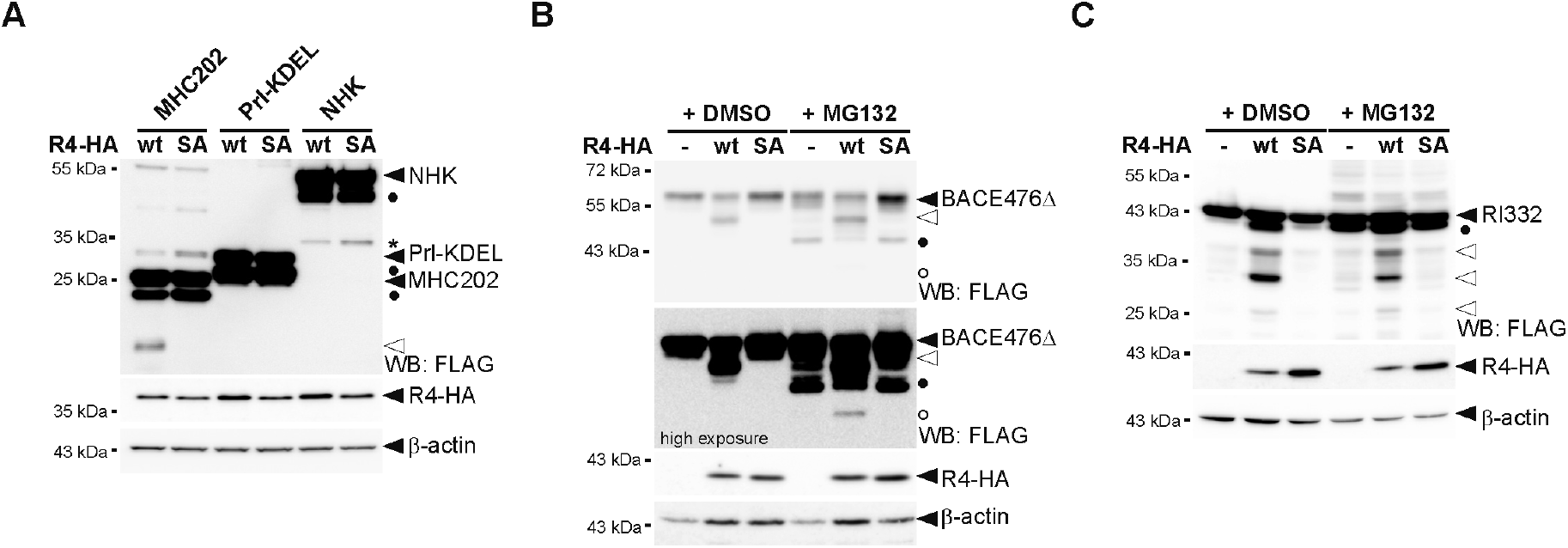
RHBDL4 cleaves several but not all soluble ERAD-L substrates. **(A)** Cleavage of MHC202 is specific, as RHBDL4 does not lead to fragments for Prl-KDEL and NHK even when the proteasome is inhibited by MG132 (2 μM). Hek292T cells were co-transfected with N-terminally FLAG-tagged MHC202, Prl-KDEL or NHK with either HA-tagged RHBDL4 (R4-HA) wild-type (wt) or the SA mutant and analyzed by western blotting (WB). Filled triangle, full-length glycosylated proteins; open triangle, MHC202 cleavage product; asterisk, RHBDL4 independent NHK degradation intermediate; filled circle, deglycosylated full-length proteins. β-actin was used as a loading control. **(B)** Hek293T cells were co-transfected with BACE476Δ and either an empty vector (-), R4-HA wt, or the catalytic inactive SA mutant. RHBDL4 generates an N-terminal 40-kDa cleavage fragment (open triangle) that is degraded by the proteasome as shown by increased steady-state level upon MG132 treatment (2 μM) compared to vehicle control (DMSO). In the presence of MG132 (2 μM), the 34-kDa deglycosylated full-length BACE476Δ (filled circle) and traces of a deglycosylated form of the RHBDL4-generated cleavage fragment (open circle) become visible. β-actin was used as a loading control. **(C)** Cleavage assay as in (B), but with N-terminally FLAG-tagged RI332 as substrate showing cleavage fragments in the range of 25 to 35 kDa (open triangles). Filled circle, deglycosylated full-length RI332. β-actin was used as a loading control. Data information: For clarity, for panels A-C representative experiments of 3 independent replicates are shown.

Taken together, these results show that in addition to unstable membrane-integral proteins (Fleig et al., 2012; Paschkowsky et al., 2018), RHBDL4 can selectively cleave some ERAD-L substrates.

### Different determinants can trigger RHBDL4-catalyzed processing

Next, we asked what requirements a protein has to fulfil to be recognized by RHBDL4. Since FLAG-tagged full-length MHC class I heavy chain (MHC-FL, Figure 4A) was not cleaved by RHBDL4 (Figure 4B), we asked whether triggering substrate ubiquitination would make it prone for cleavage. Therefore, we took advantage of the fact that as part of an immune evasion strategy, the human cytomegalovirus (HCMV) protein US11 targets MHC-FL towards ERAD E3 ubiquitin ligases (Wiertz et al., 1996). However, even though US11 prompted a higher turnover of MHC-FL (Figure 4 – figure supplement 1A), co-expression of RHBDL4 did not lead to any proteolytic processing by RHBDL4 (Figure 4B). This shows that specific substrate features and not the general ubiquitination status and turnover rate determine recognition by RHBDL4. As previous work demonstrated that a TM degron is sufficient to induce RHBDL4-catalyzed cleavage (Fleig et al., 2012), we fused the luminal part of MHC to the TM domain and cytosolic tail of a known RHBDL4 substrate, the α-chain of pre-T cell receptor (pTα). Consistent with previous findings, the TM degron was sufficient for RHBDL4-recognition (Fleig et al., 2012), leading to efficient processing of the MHC-pTα fusion protein (Figure 4C). In addition to two major cleavage sites in the context of the TM region, we observed an 18-kDa fragment in the range of the MHC202 cleavage product.

**Figure 4.**
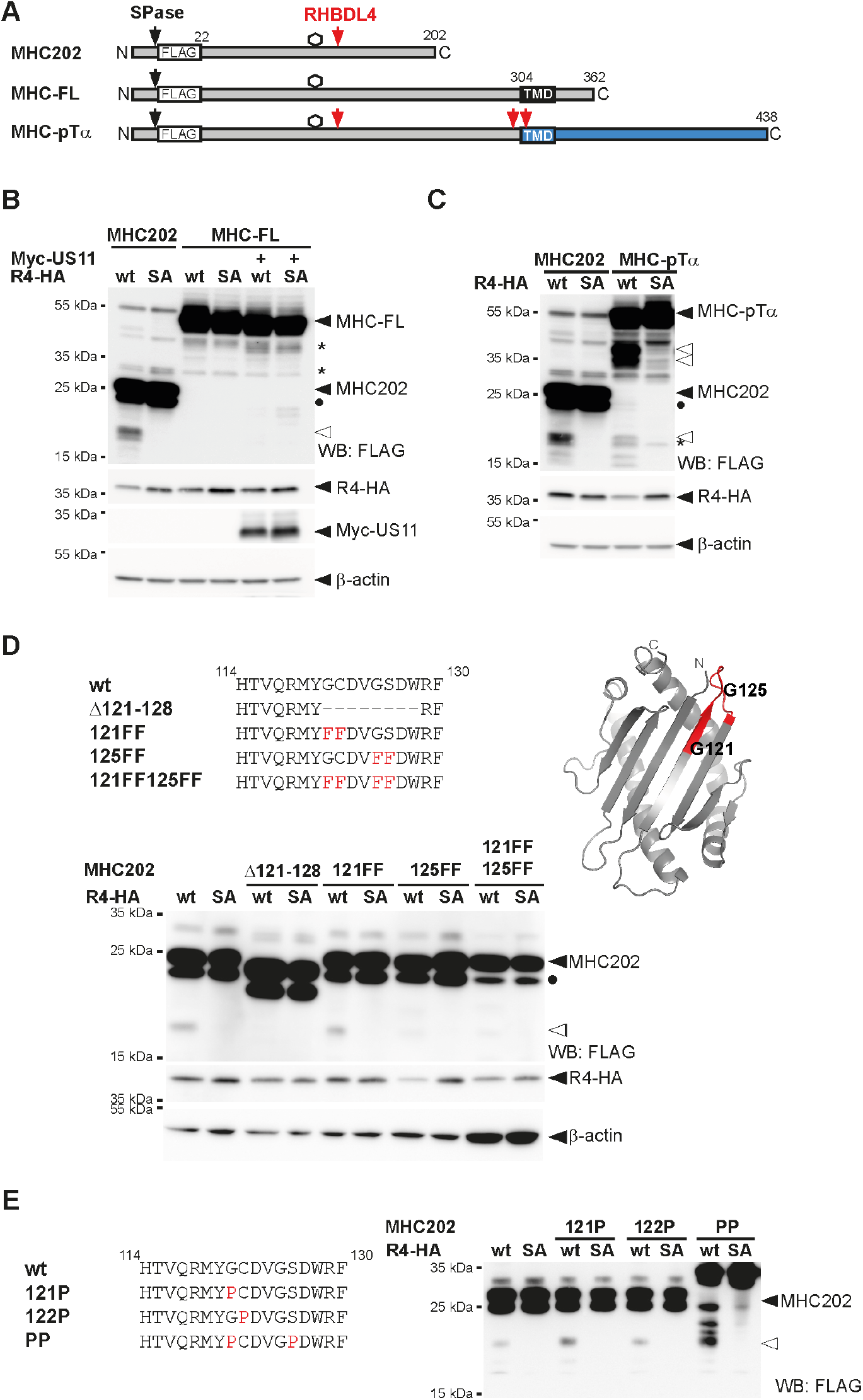
Processing by RHBDL4 is determined by specific features and not general substrate ubiquitination. **(A)** Outline of MHC202, MHC-FL and a chimaera of MHC and pTα (indicated in blue). SPase, signal peptidase; TMD, transmembrane domain. **(B)** Hek293T cells were co-transfected with N-terminally FLAG-tagged MHC202 or MHC-FL and either HA-tagged RHBDL4 (R4-HA) wild-type (wt) or SA mutant, as well as Myc-tagged US11 in the presence of MG132 (2 μM). The N-terminal cleavage fragment of MHC202 is observed in the presence of R4-HA but not for MHC-FL. Open triangle, N-terminal MHC202 fragment; filled circle, deglycosylated form of MHC202; asterisk, RHBDL4 independent degradation intermediate. β-actin was used as a loading control. WB, western blotting. **(C)** Fusion to the pTα TMD degron renders MHC susceptible for RHBDL4 cleavage in the cell-based assay as in (B). **(D)** RHBDL4 cleavage assays for the indicated MHC202 deletion and point mutants were performed in the presence of MG132 (2 μM). Cleavage by RHBDL4 occurred between amino acid 121 and amino acid 128. Small amino acids within this stretch were mutated to phenylalanine (F). Open triangle, N-terminal fragment; filled circle, deglycosylated full-length MHC202. β-actin was used as a loading control. The right panel shows the position of the two critical glycine residues (G121 and G125) and the 121-128 cleavage site region (red) in the MHC202 structure model, as shown in Figure 1A. **(E)** Cleavage assay as in (D) but with MHC202 proline point mutants. Filled triangle, glycosylated full-lenght MHC202; open triangle, N-terminal cleavage fragment of MHC202. Data information: For panels B-E representative experiments of 3 independent replicates are shown.

These results show that the C-terminal truncation of MHC202 is not strictly required for RHBDL4-catalyzed cleavage, and different determinants can lead to the same productive interaction with RHBDL4.

### RHBDL4 cleaves at a defined site, but additional features determine substrate selection

Processing of the membrane-anchored MHC-pTα in the same region as MHC202 supports the notion that RHBDL4 preferentially cleaves at specific amino acid residues. For bacterial rhomboid proteases, a loose consensus sequence with small side chains at the scissile peptide bond has been shown to at least partially determine cleavage specificity (Strisovsky et al., 2009). Hence, we narrowed down the site of RHBDL4-catalyzed cleavage and then mutated small amino acids within this stretch to phenylalanines. For MHC202, cleavage by ectopically expressed RHBDL4 was abolished in a mutant with a deletion between amino acid 121 and amino acid 128 (Figure 4D). Within this stretch, four small amino acids are found in two pairs, namely glycine-121 (G121), cysteine-122 (C122), glycine-125 (G125) and serine-126 (S126). Only mutation of all four residues to phenylalanine (121FF,125FF) abolished cleavage completely, whereas mutating the second pair (125FF) partially reduced cleavage (Figure 4D). This result indicates that the major processing occurs at G125, but G121 provides an alternative cleavage site. Interestingly, G125 is located at a surface-exposed loop between two antiparallel β-sheets forming the hydrophobic interface of the α1-α2-domains to the juxtamembrane α3-domain in full-length MHC, the latter which is deleted in MHC202 (Figure 1A and 4D). Of note, mutation of small residues in the MHC202 cleavage site region to proline, which for bacterial rhomboids has been shown to prevent the processing of the nearby peptide bond (Strisovsky et al., 2009), increased RHBDL4- catalyzed cleavage (Figure 4E). This was particularly pronounced in the glycine-121-proline, serine-126-proline double mutant (PP). For this mutant, which due to its unfolded state shows an apparent higher molecular weight on SDS-PAGE, at least three additional RHBDL4-induced cleavage products are detectable (Figure 4E). Since proline is precited to break secondary structure elements, these results indicate that the cleavage site accessibility has a major impact on MHC202 processing by RHBDL4. Overall, we provide evidence that RHBDL4 substrate selection is a multi-layer process with sequence-specific recognition of the scissile peptide bond contributing to specificity, but the secondary structure and the overall protein stability playing a dominating role.

### The erlin ERAD complex interacts with RHBDL4 and MHC202

As RHBDL4 did not primarily rely on the amino acid sequence for substrate selection, we wondered whether RHBDL4 assembles with other ERAD factors contributing to substrate recruitment. A critical step in analyzing membrane protein complexes is to combine efficient one-step affinity purification of proteins expressed at physiological levels. Therefore, we endogenously tagged RHBDL4 in Hek293T cells at its C-terminus with a single FLAG-tag using CRISPR/Cas12-mediated gene editing (Figure 5 – figure supplement 1A and B) (Fueller et al., 2020). Hek293T cells expressing FLAG-tagged RHBDL4 were grown in medium supplemented with ’heavy’ labelled amino acids, whereas the parenteral Hek293T cells were cultured in normal medium. Subsequently, the same number of cells were mixed, RHBDL4-FLAG was isolated from Triton X-100 solubilized microsomes, and co-purified interaction partners were identified by mass spectrometry (Figure 5A). The previously identified RHBDL4 cofactor p97 (Fleig et al., 2012) was 1.4-fold enriched, demonstrating the efficiency of this workflow. To identify core components of RHBDL4-dependent ERAD, we focused on proteins identified in all three replicates. Among the 20 proteins that showed enrichment in the RHBDL4-FLAG fraction greater than 1.4-fold were the chaperones BiP and calreticulin, two protein disulfide isomerases, namely PDI and Erp44, and both subunits of the regulatory glucosidase II (Supplementary file 2). Furthermore, a pair of two homologous membrane-integral ERAD factors, namely Erlin1 and Erlin2, were enriched by 1.5-fold. We reasoned that the luminal quality control factors are likely co-purified with bound RHBDL4 substrates. With a focus on the erlins, we asked whether they are part of a functional membrane protein complex. Consistent with a stable assembly, co-immunoprecipitation and western blotting confirmed co-purification of RHBDL4 with Erlin1 and Erlin2 (Figure 5B and Figure 5 – figure supplement 1C-E). Erlin1 and Erlin2 were previously demonstrated to form a MDa-ERAD complex that among other clients is involved in the degradation of the IP(3) receptor (Huber et al., 2013; Inoue and Tsai, 2017; Pearce et al., 2009), suggesting that RHBDL4 functionally interacts with this ERAD sub-branch. The E3 ligase RNF170 previously shown to interact with the erlin complex was also co-purified with ectopically expressed RHBDL4 (Figure 5 – figure supplement 1F). Interestingly, Erlin2 showed stronger interaction with RHBDL4-GFP wt than the catalytic inactive SA mutant (Figure 5 – figure supplement 1D). This result suggests that Erlin2 is not trapped by the SA mutant as would be the case for an RHBDL4 substrate. Hence, we may speculate that erlins play a role in substrate recruitment. As a putative substrate adaptor they may bind to RHBDL4 also in absence of a bound substrate but potentially dissociate from a trapped, stalled rhomboid-substrate complex. In accordance with a functional interplay of RHBDL4 with the erlin complex, blue native polyacrylamide electrophoresis (BN-PAGE) of immunoisolated FLAG-tagged RHBDL4, both endogenously and ectopically expressed, showed distinct complexes in the range of 250 kDa to 1.2 MDa, with Erlin2 co-purifying and co-migrating with the largest assembly (Figure 5C and Figure 5 – figure supplement 1G).

**Figure 5.**
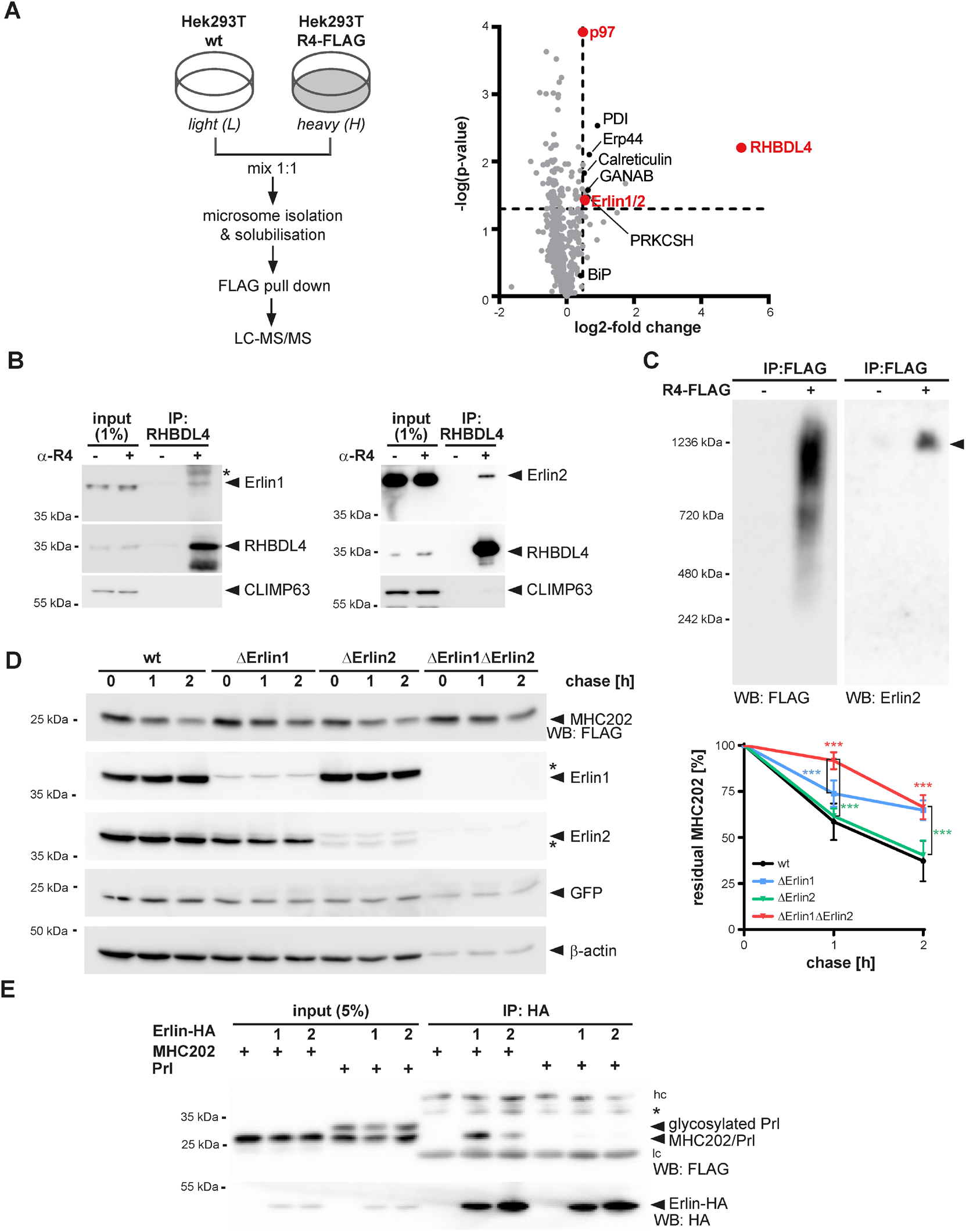
The erlin ERAD complex interacts with RHBDL4 and MHC202. **(A)** SILAC-based mass spectrometry analysis of RHBDL4 interactome from Triton X-100-solubilized microsomes obtained from Hek293T cells with chromosomally FLAG-tagged RHBDL4 (Hek293T R4-FLAG) was performed as outlined. The right panel shows a volcano plot representation of potential RHBDL4 interaction partners identified in all three replicates. **(B)** Microsome fractions of Hek293T cells were solubilized with 1% Triton X-100 and endogenous RHBDL4 was isolated by immunoprecipitation (IP). Western blotting (WB) identifies co-purification of endogenous Erlin1 and Erlin2. CLIMP63 was used as a negative control. Asterix indicates an unspecifc band. **(C)** RHBDL4 is part of an MDa-sized erlin complex. Hek293T cells transfected with empty vector (-) or FLAG-tagged RHBDL4 (R4-FLAG) were solubilized with 1% Triton X-100, immunoprecipitated for FLAG, eluted with FLAG peptides and analyzed by BN-PAGE. RHBDL4-FLAG formed several higher molecular weight complexes in addition to the 1.2 MDa complex containing Erlin2 (filled triangle). **(D)** MHC202 degradation is delayed in Erlin1 Hek293T knockout cells (ΔErlin1) compared to wild-type Hek293T cells (wt), as shown by cycloheximide (CHX) chase. To block potential compensation by ER-phagy, cells were pretreated with 100 nM BafA1 for 3h. In Erlin1/Erlin2 Hek293T double knockout cells (ΔErlin1ΔErlin2), MHC202 is significantly stabilized compared to parental Hek293T cells. To ensure homogenous expression of MHC202 within each cell line, GFP expressed from a downsteam internal ribosome entry site (IRES) and endogenous β-actin were used as controls. Asterisks indicate cross-reacting Erlin1 and Erlin2 signals. WB quantification of four independent experiments is shown in the right panel (means ± SEM, n=4, ∗∗∗p ≤ 0.001 (two-way ANOVA)). **(E)** HA-tagged Erlin1 (1) and Erlin2- (2) specifically interact with MHC202 but not with prolactin (Prl). Hc, heavy chain; lc, light chain; asterisk, unspecific band. Data information: For clarity, for panels B-E representative experiments of 3-4 independent replicates are shown.

To test our hypothesis that erlins are substrate-adaptors for RHBDL4, we generated single and double Erlin1 and -2 knockout cells and tested the stability of MHC202 by cycloheximide chase. In order to block compensation by lysosome-based pathway such as ER-phagy (Molinari, 2021), we treated cells with the vacuolar ATPase-inhibitor bafilomycin A1 (BafA1) (Figure 5 – figure supplement 1J). Interestingly, knockout of Erlin1 leads to a reduced turnover of MHC202, whereas degradation of MHC202 was not significantly delayed upon Erlin2 knockout (Figure 5D and Figure 5 – figure supplement 1H and I). Since the erlins share high sequence similarity (Pearce et al., 2009), we hypothesized that knockout of one erlin protein could be compensated by the other one. Indeed, double knockout of Erlin1 and Erlin2 further slowed down the degradation of MHC202 (Figure 5D). Consistent with a direct role in recognizing RHBDL4 substrates, immunoprecipitation of Erlin1-HA or Erlin2-HA pulled down FLAG-tagged MHC202 but not the stable, secreted control protein Prl (Figure 5E).

Altogether, these results show that RHBDL4 forms a MDa-complex with both Erlin1 and Erlin2 and that both erlins engage with the RHBDL4 substrate MHC202.

### RHBDL4 facilitates the removal of aggregation-prone ERAD-L substrates

In addition to canonical ERAD, Erlin2 was shown to act as a chaperone on the artificially designed, ER-targeted protein termed ER-beta (ERβ) (Figure 6 – figure supplement 1A), which like MHC202 is aggregation-prone (Vincenz-Donnelly et al., 2018). As Erlin2 and RHBDL4 are part of one complex, we wondered whether RHBDL4 also interacts with and degrades ERβ. Indeed, the catalytic inactive SA mutant of RHBDL4 traps ERβ resulting in co-immunoprecipitation of ERβ with RHBDL4-SA but not wt (Figure 6A). This mirrors the behaviour of RHBDL4 substrates like MHC202 (Figure 1E) or pTα (Fleig et al., 2012).

**Figure 6.**
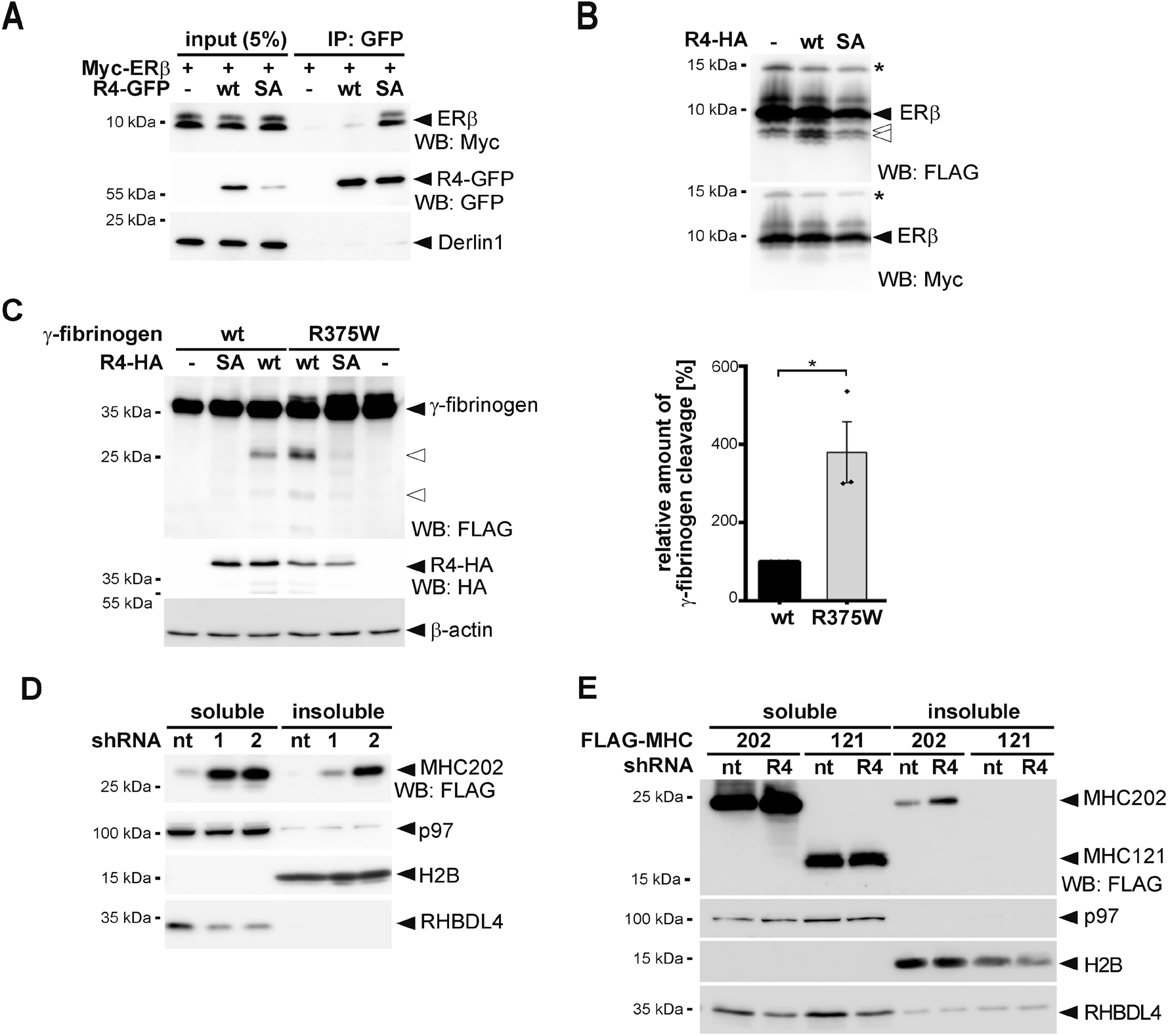
RHBDL4 reduces the burden of aggregation-prone ERAD-L substrates. **(A)** The aggregation-prone model protein ER-beta (ERβ) harbouring an N-terminal Myc tag interacts with the catalytic SA mutant of GFP-tagged RHBDL4 (R4-GFP) as shown by immunoprecipitation (IP), whereas no interaction is observed for the wild-type (wt) construct. WB, western blotting. **(B)** ERβ harbouring an N-terminal Myc-tag and a C-terminal FLAG-tag was co-expressed with HA-tagged RHBDL4 (R4-HA) as indicated. Tris-bicine-urea SDS PAGE and western blot (WB) analysis reveal at least two C-terminal cleavage fragments (open arrows) along with full-length ERβ and an undetermined post-translational modification (star). **(C)** A mutation in fibrinogen α-chain (R375W) that increases the aggregation propensity also increased generation of two N-terminal fragments (open arrows) by ectopically expressed HA-tagged RHBDL4 (R4-HA) in Hek293T cells. β-actin was used as a loading control. Western blot quantification of three independent experiments is shown in the right panel (means ± SEM, n=3, ∗p ≤ 0.05 (Student’s t-test)). **(D)** MHC202 steady-state levels were analysed in Hek293T cells transfected with two independent shRNAs targeting RHBDL4 (R4-1 and R4-2) or non-targeting control (nt) followed by NP40 lysis and WB analysis of the soluble and detergent-insoluble fraction. p97 was used as a loading control for the soluble fraction and H2B for the insoluble fraction. **(E)** MHC121 mimicking the RHBDL4-generated N-terminal cleavage fragment was not recovered in the NP40 insoluble fraction, in contrast to MHC202, upon RHBDL4 shRNA knockdown (R4) compared to non-targeting control (nt) in Hrd1 knockout cells. p97 was used as a loading control for the soluble fraction and H2B for the insoluble fraction. Data information: For clarity, for panels A-E representative experiments of three independent replicates are shown.

Consistent with this, knockdown of RHBDL4 increased the ERβ steady-state level (Figure 6 – figure supplement 1B). Furthermore, co-expression of RHBDL4 wt with ERβ increases the generation of a C-terminal cleavage fragment (Figure 6B). This raised the question of whether RHBDL4 might be of general importance for the turnover of aggregation-prone peptides. Interestingly, the disease-associated, aggregation-prone Aguadilla variant of the fibrinogen γ-chain harbouring the arginine-375-tryptophane (R375W) substitution (Brennan et al., 2002; Kruse et al., 2006) was cleaved four times more compared to the γ-chain wt (Figure 6C ). This indicates that, despite an almost unchanged amino acid sequence, the biophysical property of an aggregation-prone ERAD-L substrate likely targets γ-fibrinogen into the RHBDL4-dependent ERAD clipping pathway. Consistent with this, knock out of Erlin1 and Erlin2 leads to a significant increase of the steady-state level of γ-fibrinogen-R375W (Figure 6 – figure supplement 1B).

In order to determine the impact of RHBDL4 on clearance of aggregation-prone species of ERAD-L substrates, we analyzed the Nonidet P40 (NP40)-insoluble fraction by western blotting (Valetti et al., 1991). In addition to increasing the steady-state level of MHC202, knockdown of RHBDL4 also increases the level of MHC202 recovered in the NP40 insoluble fraction (Figure 6D) but not for a truncated version of MHC202 (MHC121), corresponding to the RHBDL4-generated N-terminal cleavage fragment. MHC121 is not recovered in the NP40 insoluble fraction even upon RHBDL4 knockdown (Figure 6E). Of note, this further truncated version of MHC121 is rapidy degraded so that we used Hrd1 knockout cells for this assay. Consistent increase of NP40 insoluble protein aggregates upon knockdown of RHBDL4 were also observed for the aggregation-prone ERAD-L substrates ERβ, γ-fibrinogen wt and the R375W mutant (Figure 6 – figure supplement 1C-E). Taken together, these results indicate that RHBDL4-catalyzed cleavage prevents self-aggregation of MHC202 and other ERAD-L substrates. The molecular mechanism of how the RHBDL4-erlin complex recognizes aggregation-prone protein conformations and how RHBDL4-catalyzed clipping facilitates dislocation into the cytoplasm are essential questions that remain to be solved in the future.

## Discussion

Protein aggregation in cells is an abnormal condition associated with ageing and human disorders ranging from diabetes to neurodegeneration (Labbadia and Morimoto, 2015; Reis-Rodrigues et al., 2012). While multiple safeguards are known to cope with cytoplasmic protein aggregates (Koga et al., 2011; Mogk et al., 2018), little is known about pathways that clear aggregating proteins from the ER lumen. Our results show that the rhomboid protease RHBDL4 contributes to the turnover of soluble, aggregation-prone ERAD-L substrates. While this substrate class is commonly degraded through a Sel1, Hrd1, and derlin-dependent retrotranslocation route (Christianson and Ye, 2014; Ruggiano et al., 2014), aggregation-prone conformations of the same substrates may be targeted by the erlin complex to RHBDL4 for cleavage (Figure 7). We suggest that this rhomboid-catalyzed clipping mechanism may facilitate protein turnover by generating shorter fragments that are more easily dislocated into the cytoplasm for proteasomal degradation. Under conditions when RHBDL4-dependent ERAD is compromised, or the substrate-load exceeds its capacity, various ERAD-L substrates aggregate, highlighting the importance of this proteostasis mechanism.

**Figure 7.**
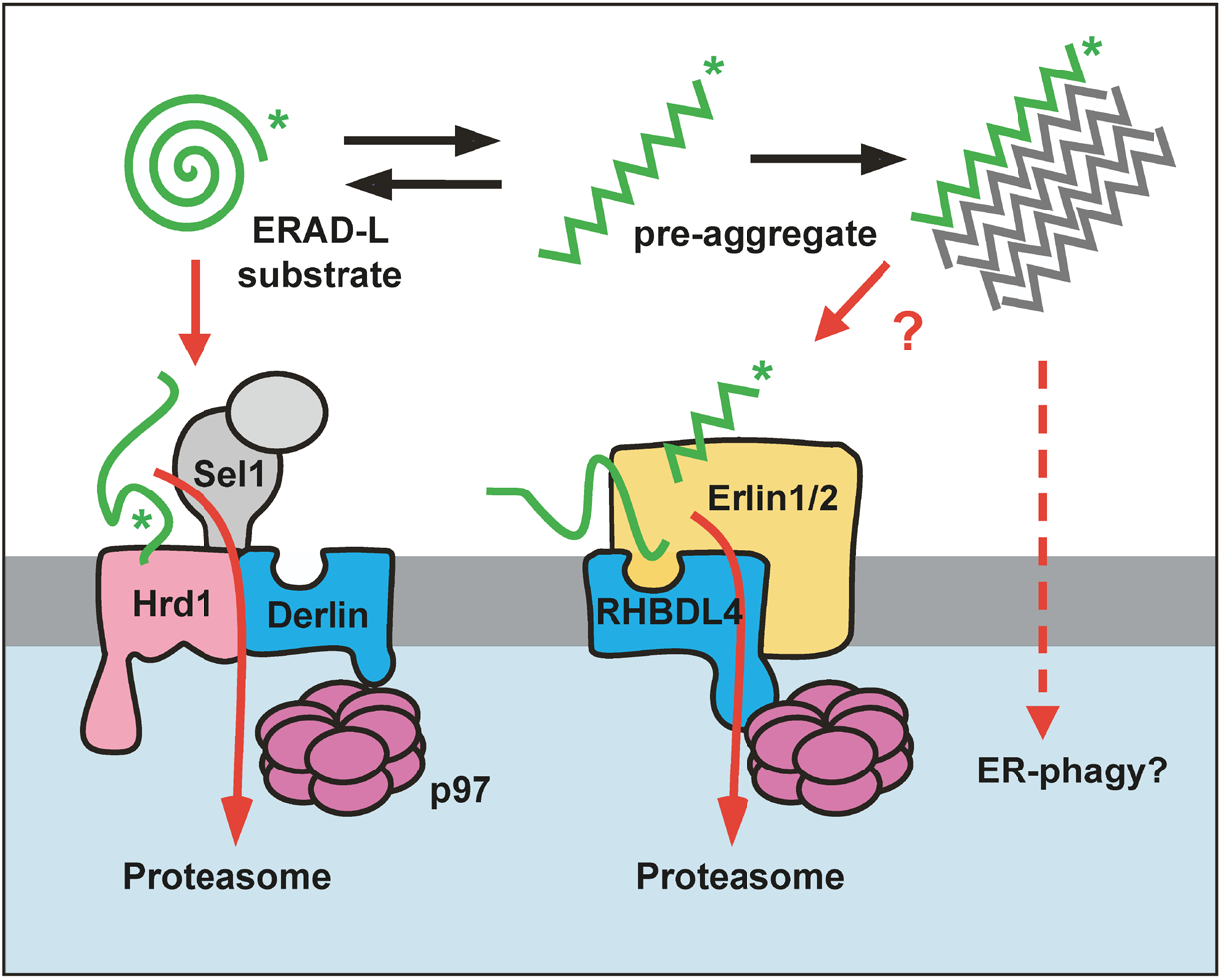
Model for RHBDL4-erlin-mediated clearance of pre-aggregates. Monomeric ERAD-L substrates are predominantly degraded through the canonical Sel1, Hrd1 and derlin-dependent retrotranslocation pathway, whereas the erlin complex targets aggregation-prone conformations to RHBDL4. RHBDL4-catalyzed clipping facilitates retrotranslocation of cleavage fragments in a process that is reminiscent to the derlin-induced recognition by Hrd1 and derlins. Upon increasing their concentration, pre-aggregated ERAD-L substrates form oligomers that may become disassembles and presented for RHBDL4-catalyzed cleavage by erlins. Large, macroscopic aggregates cannot be targeted to the ERAD pathway and may become subject to ER-phagy.

### RHBDL4 binds erlins to target aggregation-prone ERAD-L substrates for degradation

Biochemical analysis suggests that rhomboid proteases do not need any invariant subunit and may act as single-chain proteases (Lemberg et al., 2005; Urban and Wolfe, 2005). This is a striking difference to the aspartic intramembrane protease presenilin, which in order to become active has to assemble with three invariant subunits, Nicastrin, PEN2 and APH1, forming the γ-secretase complex (Edbauer et al., 2003; Kimberly et al., 2003; Takasugi et al., 2003). Here, we reveal by shotgun proteomics of genomically tagged RHBDL4 and validating by western blot analysis that the two putative substrate recruitment factors Erlin1 and Erlin2 (Pearce et al., 2007; Pearce et al., 2009) are in a native complex with RHBDL4. A previous proximity proteomics approach did not reveal significant interaction of RHBDL4 to membrane integral components (Ikeda and Freeman, 2019). However, the study by Ikeda and Freeman used a BirA fused to RHBDL4’s C-terminal tail that likely does not get in proximity to erlins that lack any prominent cytoplasmic portion (Pearce et al., 2009). In the light of our analysis of the native RHBDL4 interactome and a recent study on the mitochondrial rhomboid protease PARL (Wai et al., 2016), we may now speculate that also rhomboids form higher-order assemblies. The RHBDL4 complex shows a striking parallel to another intramembrane protease involved in ERAD: the aspartic protease SPP (signal peptide peptidase), which also forms several higher-order assemblies with specific ERAD components (Chen et al., 2014; Stagg et al., 2009). Our BN-PAGE analysis revealed several RHBDL4 containing complexes after solubilization with Triton X-100. This includes an assembly >1 MDa containing endogenous Erlin2. Previous work has shown the interaction of erlins with ERAD substrates as diverse as the IP(3) receptor (Lu et al., 2011; Pearce et al., 2007; Pearce et al., 2009) and the artificially designed aggregation-prone luminal peptide ERβ (Vincenz-Donnelly et al., 2018). Intriguingly, the erlin complex is predicted to form an assembly similar to chaperonins, albeit without ATPase activity and was hypothesized to bind hydrophobic stretches that are a hallmark for aggregating proteins (Pearce et al., 2009). The interplay of erlin-mediated recognition and RHBDL4-catalyzed clipping may help to lower protein aggregation in the ER lumen (Vincenz-Donnelly et al., 2018). While globular misfolded proteins are primarily targeted to the canonical Hrd1 pathway (Christianson and Ye, 2014; Ruggiano et al., 2014), aggregation-prone peptide conformations may be recognized by targeted towards the RHBDL4-erlin complex (Figure 7). Interestingly, erlins are members of the SPFH (stomatin, prohibitin, flotillin, HfC/K) family that also include Stomatin-like protein 2 (SLP2) (Browman et al., 2007; Browman et al., 2006). SLP2, in turn, was shown to assemble in a 2-MDa complex with the mitochondrial rhomboid protease PARL and the *i*-AAA-protease YME1L, where SLP2 is thought to regulate PARL-catalyzed intramembrane proteolysis (Wai et al., 2016).

Despite this striking similarity, it remains to be seen whether erlins comparably control RHBDL4 activity. Considering that prohibitins, the closest relatives of erlins, form higher molecular weight complexes regulating *m*-AAA proteases (Steglich et al., 1999) this could likely be a mechanism shared by several proteases linked to proteostasis control.

### Recognition of ERAD-L substrates by the membrane-integral rhomboid active site

The crystal structures of the *Escherichia coli* rhomboid protease GlpG revealed the active site to be located several Ångstroms beneath the membrane surface, in the centre of a six TM helix-bundle (Wang et al., 2006). A combination of structural and biochemical studies on bacterial rhomboids provided evidence for a lateral lipid-embedded substrate gate and a surface-exposed active site opening, which is temporally shielded by a flexible loop structure (for review see (Lemberg and Strisovsky, 2021)). While helical, lipid-embedded substrate TM segments are thought to unfold into the active site via the membrane-embedded lateral gate (Cho et al., 2016), it is conceivable that for RHBDL4, the surface-active site opening allows ERAD-L substrates lacking any TM anchor to enter the active site from the ER lumen. In a related manner, we and others observed rhomboid cleavage within ectodomains and loops of membrane proteins (Fleig et al., 2012; Knopf et al., 2020; Maegawa et al., 2007; Paschkowsky et al., 2018) and *in vitro* detergent-solubilized rhomboids are known to cleave soluble model substrates (Arutyunova et al., 2018; Wang et al., 2006). Overall, at least two different substrate recognition routes emerge for RHBDL4: one for membrane proteins and one for soluble ERAD-L substrates, which both lead to clipping and subsequent degradation by the proteasome.

### Rhomboid-fold as a conserved feature in retrotranslocation

The here observed cleavage of a soluble ERAD-L substrate may be an analogue to the interaction of derlins with ERAD-L substrates during Hrd1-mediated retrotranslocation (Wu et al., 2020). While Der1 in yeast is specific for soluble substrates (Carvalho et al., 2006; Denic et al., 2006) and a second derlin Dfm1 only deals with membrane proteins (Neal et al., 2018), RHBDL4 and mammalian derlins may act on both membrane-integral and soluble substrates. Although the exact mechanism of retrotranslocation remains to be determined, recent cryo-EM structures of ERAD complexes have revealed first insights (for review see (Lemberg and Strisovsky, 2021). Most prominently, a structural model of the yeast Hrd1 complex indicates that the ERAD-L substrates are inserted into the plane of the membrane via the rhomboid fold of Der1 and pass the lipid bilayer in between two half-channels formed by Hrd1 and Der1, respectively (Wu et al., 2020). Molecular dynamics simulation and comparison to previous work on the bacterial rhomboid GlpG suggest that this energetically unfavoured event is facilitated by a lipid thinning effect induced by both, Hrd1 and Der1. Our observation that several RHBDL4 mutants stabilize uncleaved substrates while they are looped into the cytoplasm indicates that also RHBDL4 contributes to retrotranslocation. Strikingly, mutations of the conserved GxxxG motif in human Derlin-1 revealed a very similar dislocation intermediate (Greenblatt et al., 2011). For RHBDL4 the default pathway appears to be substrate clipping and retrotranslocation of cleavage fragments. However we hypothesize that based on its homology to Der1, in concert with other ERAD factors RHBDL4 may also contribute in a non-proteolytic manner to protein dislocation into the cytoplasm. Similarly, the bacterial rhomboid protease YqgP has both a proteolytic function and acts as a pseudorotease when it recruits conformational variants of a membrane transporter to the AAA-protease FtsH for degradation (Began et al., 2020). The parallel of bacterial rhomboid proteases in membrane protein quality control (Began et al., 2020; Liu et al., 2020) to derlins and RHBDL4 in ERAD suggests that rhomboid family proteins represent an ancient proteostasis factor (Knopf and Lemberg, 2020). While initially evolved as proteases, certain rhomboids like the derlins may have lost their catalytic activity during eukaryotic evolution.

We may speculate that they retained their role in recognising aberrant proteins, but the ubiquitin-proteasome system took over the degradation function. Hence, RHBDL4, with its serine intramembrane protease active site, may be seen as an ancestral form still combining the protease and pseudoprotease mechanism.

### A role of RHBDL4 in aggregate removal

Aggregates are higher molecular structures commonly no longer soluble in nonionic detergents (Valetti et al., 1991). Seen from this angle, the increase of NP40-insoluble MHC202 under RHBDL4 knockdown first shows that MHC202 tends to aggregate and second, it suggests that RHBDL4 is important for the removal of aggregation-prone proteins. The role of RHBDL4 in clipping aggregation-prone ERAD-L substrates is corroborated by our finding that the Aguadilla mutant of fibrinogen γ-chain, predestined to form aggregates (Kruse et al., 2006), is cleaved four times more than the wt protein. Likewise, we observed that the aggregation-prone model protein ERβ functionally interacts with RHBDL4. Altogether, these results suggest that RHBDL4, in cooperation with the erlin complex, cleaves and thereby induces the degradation of aggregation-prone ERAD-L substrates. For the substrates analyzed within our study, this affects only a small fraction that may start aggregating with a lag phase of several hours, while the initial fast turnover is dominated by Hrd1-mediated retrotranslocation. In contrast, given the limited dimension of the Hrd1 complex (Wu et al., 2020) or any other putative alternative retrotranslocon depending on RHBDL4, macroscopic protein aggregates might be removed by ER-phagy or a vesicle-based lysosomal degradation route (Figure 7) (Fregno et al., 2018; Fu and Sztul, 2009). Hence, in addition to controlling the integrity of the membrane proteome as previously described (Fleig et al., 2012), RHBDL4 serves as an important fail-safe mechanism for ER luminal protein homeostasis by lowering the concentration of aggregation-prone luminal ERAD-L substrates.

Further insights into RHBDL4 complex composition and identification of additional endogenous substrates likely will unveil important cellular mechanisms. These insights will be indispensable to utilize the capacity of RHBDL4 in pre-aggregate removal for therapeutic application.

## Materials and Methods

### Plasmids and RNA interference

Unless otherwise stated, all constructs were cloned into pcDNA3.1+ (Invitrogen). Construct encoding human RHBDL4 with an N-terminal triple HA-tag and C-terminal GFP-tag have been described previously (Knopf et al., 2020). Constructs for HA-tagged human RHBDL4-ΔC and RHBDL4-ΔVBM were cloned by subcloning residues 1 to 268 (ΔC) and residues1 to 300 (ΔVBM), respectively (Fleig et al., 2012). For generating point mutants, a site-directed mutagenesis strategy was used. For affinity purification by immunoprecipitation and peptide elution, a C-terminal single FLAG-tagged mouse RHBDL4 was cloned (Fleig et al., 2012).

Plasmids encoding triple FLAG-tagged RI332, secreted human prolactin and Prl-KDEL were described previously (Fleig et al., 2012). A truncated 202-amino acid long version of human MHC class I heavy chain A2 (UniProt ID O78126) with a C-terminal FLAG-tag was cloned into pCMV-S11 (Sandia BioTech). N-terminal triple FLAG-tagged versions of MHC-FL, MHC202 (comprising residues 21 to 202 of the MHC ORF), OS9 (UniGene ID Hs. 527861, IMAGE:2964645), NHK (gift from R. Kopito), BACE476Δ (gift from M. Molinari), fibrinogen γ- chain wt and -R375W (gift from J. Brodsky) were generated by subcloning the respective open reading frames omitting their signal sequences into a pcDNA3-based expression vector containing a signal sequence fused to a triple FLAG-tag (Fleig et al., 2012). For cycloheximide experiments, the N-terminal triple FLAG-tagged version of MHC202 was subcloned into pCDH-IRES-GFP (Meissner et al., 2011). The glycosylation mutants MHC202-K197N and MHC202-N100Q-K197N were cloned with a C-terminal triple FLAG-tag followed by an S-tag. The MHC-pTα chimera was generated by overlap extension PCR, fusing residues 22-304 of MHC-FL to the TM domain and C-terminus of pTα (residues 147- 281). For stable expression, FLAG-MHC202 was subcloned into pcDNA5/FRT/TO (Invitrogen). Myc-tagged HCMV strain AD169 US11 (UniProt ID P09727) was ordered after codon optimization as gBlock (IDT) and cloned into pcDNA3.1+. Constructs encoding GFP-tagged ERdj3-GFP-3Gly (gift from M. Schuldiner) (Ast et al., 2016), FLAG-tagged RNF170, HA- and FLAG-tagged human Erlin1 and Erlin2 (gift from R. Wojcikiewicz) (Pearce et al., 2007; Pearce et al., 2009) and Myc-tagged ERβ (gift from M. Hipp) (Vincenz-Donnelly et al., 2018), the ER marker RFP-KDEL (Altan-Bonnet et al., 2006) were described previously. For cleavage assays, ERβ was cloned with an N-terminal Myc and a C-terminal triple FLAG-tag into pcDNA3.1. To rescue Hrd1 in Hrd1 knockout cells, untagged Hrd1 was generated by subcloning the respective open reading frame (UniProt ID Q86TM6-3) into a pcDNA3-based expression vector. For transient knockdown, the small hairpin (shRNA)-expressing vectors pSUPER.neo (R4-1) (Fleig et al., 2012) and a pRS vector-based construct targeting 5’- ATGAGGAGACAGCGGCTTCACAGATTCGA-3’ (R4-2) (OriGene) were used. As non-targeting (nt) control pSUPER.neo targeting 5’-ACAGCUUGAGAGAGCUUUA-3’ designed for knockdown of RHBDL4 in COS7 cells (but not human cells) was used. For generating single guide (sgRNA) target sequences for Erlin2, the E-CRISPR tool (http://www.e-crisp.org) was used (Heigwer et al., 2014). The target sequence 5’- CACCGGCTGTGCACAAGATAGAAGA-3’ was then cloned in a BbsI linearized px459.v2 vector containing puromycin selection. For the siRNA screen, an ON-TARGETplus SMARTpool custom library (Thermo Fisher Scientific) was used (Supplementary file 1).

Knocking down RHBDL4 and NGLY 25 pmol ON-TARGETplus SMARTpools human siRNA (Dharmacon) were used. The amount of transfected siRNA was kept constant within an experiment by the addition of scrambled control.

### Cell lines and transfection

Hek293T cells were cultured in DMEM (Invitrogen) complemented with 10% fetal bovine serum at 37°C in 5% CO2. Transient transfections were performed using 25 kDa linear polyethyleneimine (Polysciences) (Durocher et al., 2002) as had been described (Fleig et al., 2012). Typically, 500 ng plasmid encoding substrate candidate and 100 ng plasmid encoding RHBDL4 were used per well of a 6-well plate. Total transfected DNA (2 μg/well) was held constant by the addition of empty plasmid. If not otherwise stated, cells were harvested 48 h after transfection. For short-term knockdown, siRNA was transfected using RNAimax (Invitrogen) transfection reagent according to manufacturer recommendation. For simultaneous transfection of siRNA and plasmid DNA Lipofectamin2000 (Invitrogen) transfection reagent was applied according to manufacturer protocol. For inhibition of the proteasome or p97, approx. 32 h post-transfection either 2 μM MG132 (Calbiochem) or 2.5 μM CB-5083 (ApexBio) were added from a 10,000 x stock in dimethylsulfoxide (DMSO). As a vehicle control, the same amount of DMSO was used. Subsequently, cells were further incubated and harvested 16 h later. Cells were lysed in SDS sample buffer (see below).

To prepare doxycycline-inducible stably transfected cells, pcDNA5/FRT/TO/FLAG-MHC202, Flp-In Hek293T-REx cells were co-transfected with pOG44 (Invitrogen), followed by selection with hygromycin B (125 μg/ml). RHBDL4 knockout cells had been described previously (Knopf et al., 2020). For generating Erlin2 knockout cells, 1 μg of CRISPR/Cas9 vector were transfected into Hek293T. After 24 h, a single cell dilution was performed. Clones were analyzed by western blotting and sequencing of a PCR amplicon obtained from genomic DNA. Primers used for validation of Erlin2 knockout cells were: 5’- CTTGAGCAACGGCTGTATCC-3’ and 5’- AATCACCACCCATGGCATCAT-3’ leading to a 610 bp amplicon. Erlin1 knockout cells and Erlin1Erlin2 double knockout cells were respectively generated in parental Hek293T, and Hek293T Erlin2 knockout cells by introducing a Stop cassette in exon 3 according to previously described CRISPR/Cas12 mediated gene editing (Fueller et al., 2020). Primers used for validation were 5’- CCAGAGGTACGGTTGGTTGA-3’ and 5’-CCTTCCAAGCTTCCTGGTTCA-3’, leading to a 547 bp amplicon. Generation of chromosomally tagged RHBDL4-FLAG Hek293T cells with a single FLAG before the stop codon in the last exon by using CRISPR/Cas12 mediated gene editing has been described before (Fueller et al., 2020). Primers used for validation were: 5’- TTATGGAGCACGATGGAAGGAA-3’ and 5’-GAGATGGGAGCGTGGAAACT-3’, leading to a 634 bp amplicon. Hrd1 knockout cells were generated according to previously described CRISPR/Cas12 mediated gene editing (Fueller et al., 2020). The cells were validated by using the following primers: 5’-GGCTATTTTGCACAGCACGA-3’ and 5’- CTTCCACCTGCTCCAGAACT-3’, leading to a 786 bp amplicon. The obtained PCR amplicons were sequenced by Sanger sequencing and analyzed using CRISP-ID (Dehairs et al., 2016).

### Antibodies

The following antibodies were used: mouse monoclonal anti-FLAG (M2, Sigma), rat monoclonal anti-HA (Roche), mouse anti-myc (New England Biolabs), rabbit polyclonal anti-GFP (gift from Oliver Gruss) and mouse monoclonal anti-GFP (Roche), mouse monoclonal anti-β actin (Sigma), mouse monoclonal anti-Derlin1 (Sigma), rabbit polyclonal anti-p97 (gift from Bernd Dobberstein), rabbit polyclonal anti-H2B (Abcam), mouse monoclonal anti-Bap31 (Alexis Biochemicals), rabbit polyclonal anti-Synoviolin (Bethyl laboratories Inc.), mouse monoclonal anti-CLIMP63 (Enzo Life Sciences), rabbit polyclonal anti-Erlin1 (Sigma), rabbit polyclonal anti-Erlin2 (Sigma), rabbit polyclonal anti-LC3B (Bio Techne GmbH), rabbit polyclonal anti-RHBDL4 (Sigma) and rabbit polyclonal anti-RHBDL4 (Fleig et al., 2012).

### Trageted siRNA screen

Downregulation of the 40 candidate proteins (Figure 1B and Figure 1 – figure supplement 1A) was conducted by using two different sets of pre-designed ON-TARGETplus SMARTpool custom siRNA libraries (Thermo Fisher Scientific). p97 was used as both positive and loading control. The #1 set was tested only one time and two times for the #2 set (see additional Supplementary file 1). For quantification, MHC202 steady-state level were not normalized to loading control p97 (Figure 1 – figure supplement 1A). The knockdown efficiency of the siRNA screen was not validated.

### Microscopy

For immunofluorescence analysis, cells were either chemically fixed in PBS containing 4% paraformaldehyde for 30 min followed by permeabilization in PBS containing 0.5% Triton X- 100 for 10 min (Figure 2 – figure supplement 1D) or fixed in methanol at -20°C for 5 minutes (Figure 2 – figure supplement 1E). Subsequently, cells were washed with PBS, blocked with 20 % fetal calf serum in PBS and probed with affinity-purified anti RHBDL4 antibody (1:50; see above) and anti-FLAG antibody (1:1000). After staining with fluorescently labelled secondary antibody (Santa Cruz Biotechnology), slides were analyzed using a TCS SP5 confocal microscope (Leica).

### NP40 solubility assay

To test the influence of RHBDL4 on the solubility of proteins, 300 ng substrate expressing vector was transfected with 1000 ng shRNA and 700 ng empty vector. After 24 - 48 h of transfection, cells were pelleted and solubilized in NP40 lysis buffer (50 mM Tris-Cl, pH 7.4, 150 mM NaCl, 2 mM MgCl2, 1% Nonidet P-40) supplemented with 1xPI. After 10 min centrifugation at full speed at 4°C, supernatant corresponding to the soluble fraction was transferred into a new tube containing 4x sample buffer (see below). The pellet was dissolved in 1x sample buffer and corresponds to insoluble fraction.

### Cycloheximide chase

Cycloheximide (100 μg/ml) chase was conducted 24 h after transfection of Hek293T cells. For inhibition of the vacuolar ATPase cells were treated with 100 nM BafA1 (AdipoGen Life Sciences). Cell extracts were subjected to western blot analysis as described below.

### Pulse-chase analysis

For pulse-chase analysis, transfected Hek293T Hrd1 knockout cells were starved for 1 h in methionine/cysteine free DMEM (Invitrogen) supplemented with 10% dialysed fetal calf serum. Consequently, cells were metabolically labelled for 20 min with 55 μ Ci/ml 35S- methionine/cysteine protein labelling mix (PerkinElmer). Cells were washed with PBS and cultured in DMEM (Invitrogen) complemented with 10% fetal bovine serum. At the harvesting time point, cells were rinsed with PBS and solubilized with 1% Triton X-100 in IP buffer (50 mM HEPES-KOH, pH 7.4, 150 mM NaCl, 2 mM MgOAc2, 10% glycerol, 1 mM EGTA) followed by FLAG-IP (as described below). Samples were subjected to SDS-PAGE, and labelled proteins were visualized by a FLA-7000 phosphorimager (Fuji).

### Protease protection assay

Protease protection assay was performed using microsomes obtained by hypotonic swelling and centrifugation from Hek293T cells 24 h after transfection. To this end, cells were resuspended in isolation buffer (10 mM HEPES-KOH pH 7.4, 1.5 mM MgCl2, 10 mM KCl, 0.5 mM dithiothreitol, 10 μg/ml phenylmethylsulfonyl fluoride (PMSF)). After 10 min incubation at 4°C, cells were lysed by passing six times through a 27-gauge needle. Cellular debris and nuclei were discarded after centrifugation at 1,000 g for 5 min at 4°C. The supernatant was spun at 100,000 g for 30 min at 4°C. The membrane pellet was resuspended in rough microsome buffer (50 mM HEPES-KOH pH 7.4, 250 mM sucrose, 50 mM KOAc, 5 mM MgO(Ac)2, 1 mM dithiothreitol). Microsomal fraction was incubated with Proteinase-K (500 μg/ml) or Proteinase-K with 1% TritonX-100 for 15 min on ice. The reaction was stopped by adding 2.5 mM PMFS for 5 min on ice. Samples were resuspended in SDS sample buffer followed by SDS-PAGE and western blotting (see below).

### Immunoprecipitation and proteomics

If not indicated differently, all steps were performed at 4°C. For substrate trapping, RHBDL4- GFP expressing Hek293T cells were solubilized with 1% Triton X-100 in IP buffer (50 mM HEPES-KOH, pH 7.4, 150 mM NaCl, 2 mM MgOAc2, 10% glycerol, 1 mM EGTA), containing 1xPI and 10 μg/ml PMSF. Cell lysates were cleared by centrifugation at 10,000 g for 10 min, following pre-clearing for 1 h with BSA-coupled sepharose beads or protein A/G beads. Anti-GFP immunoprecipitation was performed using a monoclonal GFP-specific antibody in combination with protein G beads (Figure 1E) or GFP-specific single-chain antibody fragment (Rothbauer et al., 2008) coupled to NHS-activated sepharose beads (Figure 5C) as described (Fleig et al., 2012). For immunoprecipitation of HA-tagged proteins, anti-HA antibody-coupled agarose beads (Sigma) were used. For immunoprecipitation of endogenous RHBDL4, the primary antibody was added together with protein A beads for overnight incubation. For immunoprecipitation of endogenous Erlin1 or Erlin2, the primary antibody was added together with protein A beads for overnight incubation.

Immunoprecipitates were washed three times in IP buffer containing 0.1% Triton X-100 and then resuspended in SDS sample buffer followed by SDS-PAGE and western blotting (see below).

For isolation of endogenous RHBDL4 interaction partners by shotgun proteomics, Hek293T- RHBDL4-FLAG cells were grown for at least six doublings in medium supplemented with heavy amino acids (^13^C6^15^N4-L-Arg and ^13^C6^15^N2-L-Lys, from Silantes), whereas the parenteral Hek293T cells cultured in light-medium were used as control. The third replicate was performed with a label swap to minimize the experimental error. After harvesting, an equal number of cells from both cultures were mixed, and pooled microsome fraction was isolated by hypotonic swelling and centrifugation as described above. For immunoprecipitation of RHBDL4-FLAG, microsomes were solubilized with 1% Triton X-100 in IP buffer, containing 1xPI and 10 μg/ml PMSF. Cell lysates were cleared by centrifugation at 20,000 g for 10 min. Pre-clearing with protein A beads and anti-FLAG immunoprecipitation was performed as described above. The immunocomplexes were eluted in SDS sample buffer and resolved by SDS-PAGE. The lane was subdivided into three pieces, and an in-gel trypsin digest was performed. First, proteins were reduced with DTT, alkylated with iodoacetamide and then digested with trypsin. Following digestion, peptides were extracted with 50% acetonitrile/0.1% TFA and concentrated in a SpeedVac vacuum centrifuge. The sample was analyzed by a UPLC system (nanoAcquity) coupled to an ESI LTQ Oribitrap mass spectrometer (Thermo). The uninterpreted MS/MS spectra were searched against the SwissProt-human database using MaxQuant software. The algorithm was set to use trypsin as enzyme, allowing at maximum for two missed cleavage site, assuming carbamidomethyl as a fixed modification of cysteine, and oxidized methionine and deamidation of asparagines and glutamine as variable modifications. Mass tolerance was set to 4.5 ppm and 0.2 Da for MS and MS/MS, respectively. In MaxQuant the ’requantify’ and ’match between runs’ option was utilized, the target decoy method was used to determine 1% false discovery rate. All analysis was performed on the “protein groups” file using Perseus software version 1.6.5.0 (Tyanova et al., 2016) and Microsoft Excel. Label-free intensities were used to calculate the heavy over light ratios, which were averaged over all three biological replicates. P values of log2 transformed data were determined by one-sample t-test. The cutoff for a protein to be called significantly enriched was set to fold change >1.4 and p-value <0.05.

### Blue native PAGE

If not indicated differently, all steps were performed at 4°C. Hek293T cells ectopically expressing RHBDL4-FLAG or expressing chromosomally FLAG-tagged RHBDL4 were lysed with 1% Triton X-100 in BN buffer (50 mM HEPES-KOH, pH 7.4, 150 mM NaCl, 2 mM MgOAc2, 10% glycerol, 1 mM EGTA) supplemented with EDTA-free complete protease inhibitor cocktail (1xPI, Roche) and 10 μg/ml PMSF. After removing cell debris, 10 μl anti-FLAG antibody-conjugated agarose beads (M2, Sigma) were added. After a 3 h incubation, beads were washed twice with BN buffer containing 0.2% Triton X-100 and subsequently eluted with 0.5 μg/μl FLAG peptide for 30 min. A 1/40 volume of BN sample buffer (500 mM 6-aminohexanoic acid, 100 mM bis Tris pH 7.0, 5% Coomassie G250) was added before subjection onto NativePAGE Novex Bis-Tris 3-12% gradient gels (Thermo). Gels were run for 1 h at 150 V, buffer changed according to the manufacturer’s description and then continued at 230 V for 45 min. Afterwards, gels were incubated for 15 min in blotting buffer, then transferred at 85 mA for 70 min onto PVDF membrane using a tank-blotting system. The PVDF membrane was incubated in fixation solution (40% methanol, 10% acetic acid), blocked in 5% milk TBS-Tween (10 mM Tris-Cl pH 7.4, 150 mM NaCl, 0.1% Tween 20), and analyzed using enhanced chemiluminescence (see below).

### Western blotting

Transfected cells and immunoisolated proteins were solubilized in Tris-glycine SDS-PAGE sample buffer (50 mM Tris-Cl pH 6.8, 10 mM EDTA, 5% glycerol, 2% SDS, 0.01% bromphenol blue, 5% β-mercaptoethanol). All samples were incubated for 15 min at 65°C. For deglycosylation, solubilized proteins were treated with Endo H and PNGase F (New England Biolabs) according to the manufacturer’s protocol. Denaturated and fully-reduced proteins were resolved on Tris-glycine SDS-PAGE followed by western blot analysis onto PVDF membrane (Immobilon-P, 0.45 μM pore size, Merck Millipore) using enhanced chemiluminescence to detect bound antibodies (Pierce). For the analysis of ERβ-derived cleavage fragments (<10 kDa), post-nuclear supernatants from Triton X-100 solubilized cells were mixed with Tris-bicine-urea SDS-sample buffer (360 mM BisTris, 160 mM bicine, 1% SDS, 50 mM dithiothreitol, 15% sucrose, 0.01% bromphenol blue, and 0.004% Serva blue), heated at 65°C. Peptides were separated to Tris/Bicine-urea PAGE (15% T, 5% C, 8 M urea) (Wiltfang et al., 1997), transfer onto PVDF membrane with 0.2 μm pore size and analyzed by western blotting. For detection, the LAS-4000 system (Fuji) was used.

### Reproducibility and statistics

The number of biological replicates of experiments is described in the figure legends. For quantification, band intensities were measured using the Fiji ImageJ software (Schindelin et al., 2012). For statistical analysis GraphPad Prism was used. Student’s t-test was used for statistical analysis of protein steady-state levels, and two-way ANOVA was used to calculate p values of CHX chase and pulse-chase experiments..

## Data availability

The mass spectrometry proteomic data have been deposited to the ProteomeXchange Consortium (http://proteomecentral.proteomexchange.org) via the PRIDE partner repository (Perez-Riverol et al., 2019) with the dataset identifier PXD027346.

## Acknowledgements

We gratefully acknowledge the contribution of Lina Wunderle to the beginning of this project. We thank Maya Schuldiner, Richard Wojcikiewicz, Ron Kopito, Maurizio Molinari, Jeff Brodsky and Mark Hipp for reagents. We thank Matthias Feige and Sebastian Schuck for the critical reading of the manuscript. Mass spectrometry was performed at the ZMBH Core facility for mass spectrometry and proteomics. This study was supported by the Deutsche Forschungsgemeinschaft (DFG, German Research Foundation) 20134854 / SFB1036/2- TP12 (to MKL), a fellowship by the Boehringer Ingelheim Fonds (to JDK) and an intramural research program of the National Institute of Diabetes, Digestive & Kidney Diseases in the National Institutes of Health (to YY).

## Author contribution

JB and NK designed and performed most experiments and wrote the manuscript. JDK carried out experiments. JGL performed and validated the siRNA screen. NL initiated the interactome analysis and validated interaction partners. YY helped designing the project. MKL guided the project and wrote the manuscript.

## Figure Supplements Legends

**Figure 1 – figure supplement 1.**
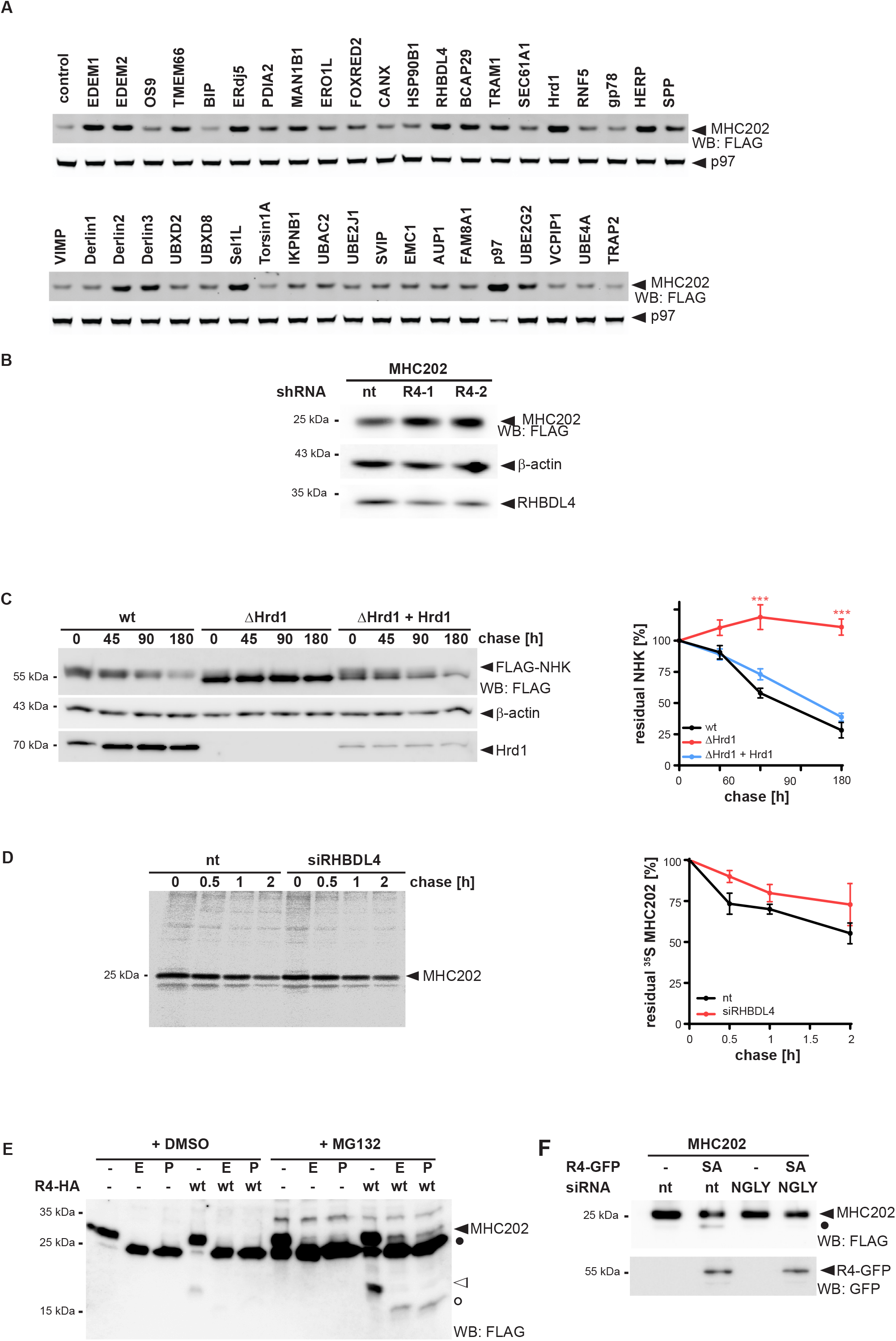
MHC202 is degraded by a concerted action of the Hrd1 complex and RHBDL4. **(A)** Influence of selected ERAD components on the steady-state level of MHC202 shown by western blot (WB) of lysates from siRNA transfected cells detecting the C-terminal FLAG tag. p97 was used as both positive and loading control. Knockdown efficiency of the siRNA screen was not determined and quantification of MHC202 steady-state level was not normalized to the loading control p97. **(B)** RHBDL4 knockdown with two independent shRNAs (R4-1 and R4-2) in Hek293T cells leads to an increase of MHC202 steady-state level when compared to a non-targeting (nt) control shRNA. β-actin was used as a loading control. **(C)** N-terminal FLAG-tagged NHK is stabilized in Hrd1 knockout cells (ΔHrd1) compared to parental Hek293T cells (wt). Co-expression of FLAG-NHK and Hrd1 rescues the degradation defect in ΔHrd1 cells. β-actin is used as a loading control. WB quantification of three independent experiments is shown in the right panel (means ± SEM, n=3, ∗∗∗p ≤ 0.001 (two-way ANOVA)). **(D)** siRNA knockdown of RHBDL4 delayed MHC202 degradation in Hek293T cells compared to the non-targeting siRNA control (nt) in metabolic pulse label chase. The right panel shows the quantification of autoradiograms of three independent experiments (means ± SEM, n=2). **(E)** MHC202 (filled triangle) and its 18 kDa N-terminal fragment (open triangle) generated by ectopically expressed HA-tagged RHBDL4 (R4-HA) are glycosylated as shown by sensitivity to Endo H (E) and PNGase (P). Filled circle, deglycosylated full-length MHC202; open circle, deglycosylated N-terminal cleavage fragment. Samples, as shown in Figure 1D, either treated with vehicle control (DMSO) or MG132 (2 μM). **(F)** Stabilization of the unglycosylated form of MHC202 (filled circle) by the catalytic inactive mutant of RHBDL4 (SA-GFP) is abolished upon knockdown of N-glycanase (NGLY) when compared to non-targeting control (nt). Data information: For clarity, for panels B-F representative experiments of three independent replicates are shown.

**Figure 2 – figure supplement 1.**
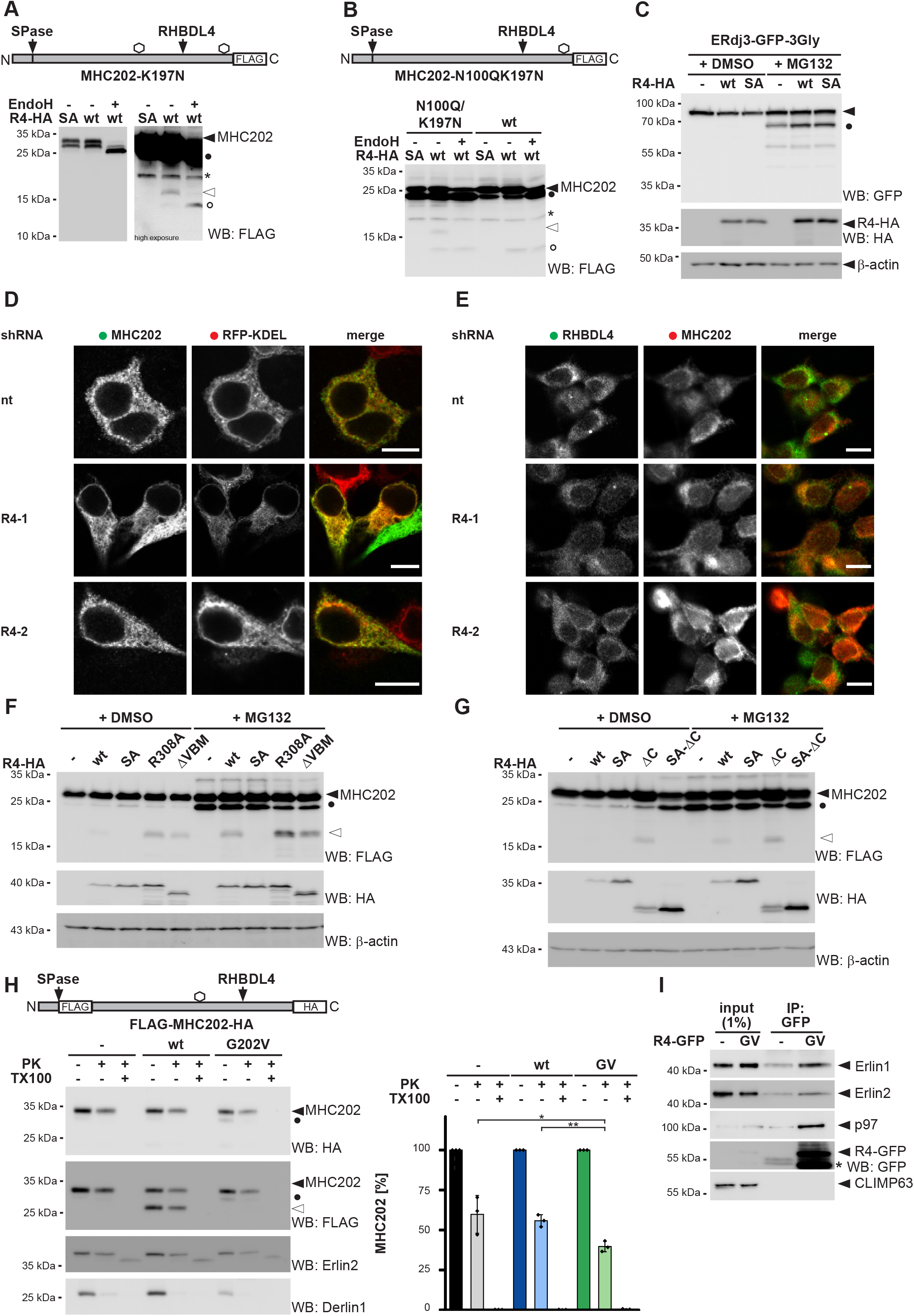
RHBDL4 knockdown leads to accumulation of MHC202 in the ER. **(A)** RHBDL4 cleaves C-terminally FLAG-tagged MHC202 with an additional C-terminal glycosylation site (K197N) post-translocational as shown by the sensitivity of the C-terminal fragment to Endo H (open triangle and open circle).Asterix, unspecific band; filled circle, deglycosylated full-length MHC202; R4-HA, HA-tagged RHBDL4; hexagon, site for N-linked glycosylation; SPase, signal peptidase; WB, western blotting. **(B)** RHBDL4 cleaves MHC202 lacking the native glycosylation site (N100Q) with a single glycosylation site (K197N) in the C-terminal portion leading to an Endo H-sensitive and a partially deglycosylated fragment (open triangle and open circle). Asterix, unspecific band; filled circle, deglycosylated full-length MHC202; hexagon, site of N-linked glycosylation; SPase, signal peptidase. **(C)** RHBDL4 does not cleave ERdj3-GFP-3Gly (filled triangle). Deglycosylated ERdj3-GFP- 3Gly (filled circle) is stabilized upon proteasomal inhibition by MG132 (2 μM). β-actin was used as a loading control. **(D)** Knockdown of RHBDL4 with two independent shRNAs (R4-1, R4-2) leads to MHC202 accumulation in the ER as shown by colocalization with RFP-KDEL; nt, non-targeting control shRNA; scale bar, 10 μm. **(E)** Knockdown of RHBDL4 with two independent shRNAs (R4-1, R4-2) increases MHC202 signal in stable T-REx Hek293T cell expressing FLAG-tagged MHC202, when compared to nt control shRNA; scale bar, 10 μm. **(F)** The N-terminal cleavage fragment (open triangle) of C-terminally FLAG-tagged MHC202 generated by the HA-tagged RHBDL4-R308A mutant is stabilized in the absence of proteasome inhibitor MG132 (2 µM). Similarly, co-expression of an RHBDL4 mutant lacking the binding motif for p97 (ΔVBM) together with MHC202 results in stabilization of the N- terminal cleavage fragment. Impaired p97 interaction of RHBDL4 leads to a slightly reduced steady-state level of deglycosylated unprocessed MHC202 form (filled circle). β-actin was used as a loading control. **(G)** HA-tagged RHBDL4 mutant lacking the C-terminal domain (ΔC) cleaves FLAG-tagged MHC202 and stabilizes the deglycosylated unprocessed form of MHC202 (filled circle) even in the absence of proteasome inhibitor MG132 (2 μM). β-actin was used as a loading control. **(H)** The accessibility of MHC202 to exogenous proteinase K (PK) was analysed in ER- derived microsomes. Hek293T cells were co-transfected with double-tagged MHC202 and either an empty vector (-), RHBDL4 wt, or the RHBDL4-G202V mutant (G202V). Hek293T- derived microsomes were incubated with PK in the presence and absence of 1 % TritonX- 100 (TX100). The R4-induced MHC202 cleavage fragment (open triangle) was protected from exogenous PK, whereas full-length MHC202 was partially accessible. Erlin2 (epitope in ER lumen) and Derlin1 (epitope in cytosol) were used as controls.Filled circle, deglycosylated full-length MHC202. HA signals of three independent experiments were quantified and shown in the right panel (means ± SEM, n=3, ∗p ≤ 0.05; ∗∗p ≤ 0.01 (Student’s t-test)). **(I)** GFP-tagged RHBDL4-G202V (R4GV-GFP) interacts with endogenous p97, Erlin1 and -2, but not CLIMP63 which was used as a negative control. Asterics indicates unspecifc band. Data information: For clarity, for all panels representative experiments of three independent replicates are shown.

**Figure 3 – figure supplement 1.**
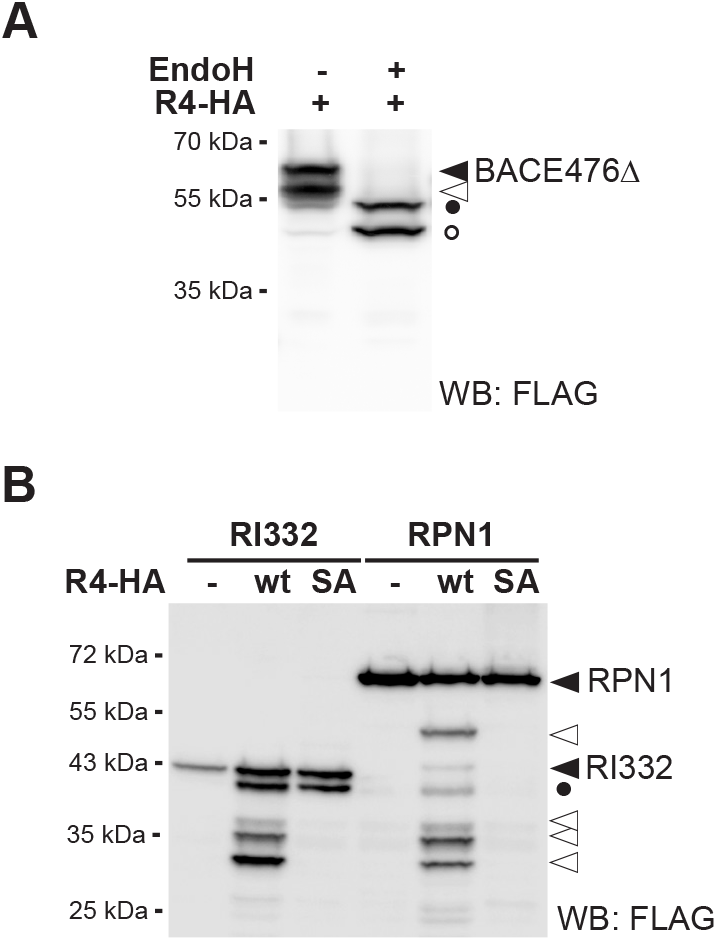
RHBDL4-catalyzed cleavage of BACE476Δ generates a glycosylated N-terminal fragment. **(A)** BACE476Δ (filled triangle) and its HA-tagged RHBDL4 (R4-HA) generated N-terminal fragment (open triangle) are glycosylated as shown by sensitivity to Endo H (filled and open circles). WB, western blotting. **(B)** Hek293T cells were co-transfected either with RI332 or RPN1 and an empty vector (-), R4-HA wild type (wt), or the catalytic inactive SA mutant. RHBDL4 generates several N-terminal cleavage fragments (open triangles). Expression of the catalytic mutant stabilizes the 40-kDa deglycosylated full-length RI332 (filled circle) even in the absence of the proteasome inhibitor MG132 Data information: For clarity, for all panels representative experiments of three independent replicates are shown.

**Figure 4 – figure supplement 1.**
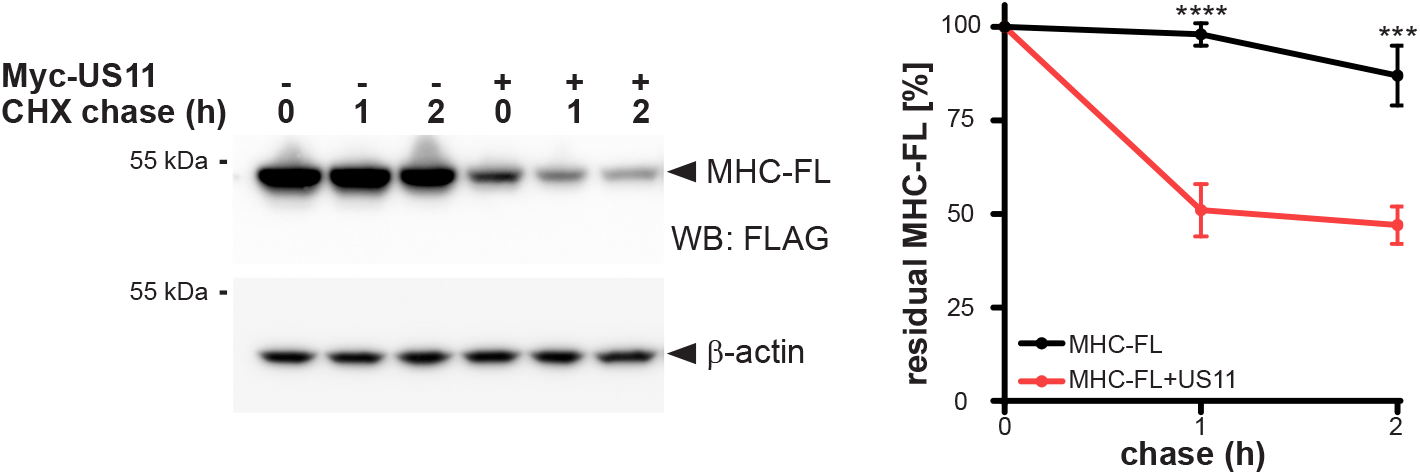
US11 increases turnover of MHC-FL. Hek293T cells were transfected with MHC-FL with or without Myc-tagged US11. 24 h post-transfection cycloheximide (CHX) was added, and cells were harvested at indicated time points (means ± SEM, n=4; ∗∗∗∗p ≤ 0.0001 (two-way ANOVA)). A representative experiment of three independent replicates is shown. WB, western blotting.

**Figure 5 – figure supplement 1.**
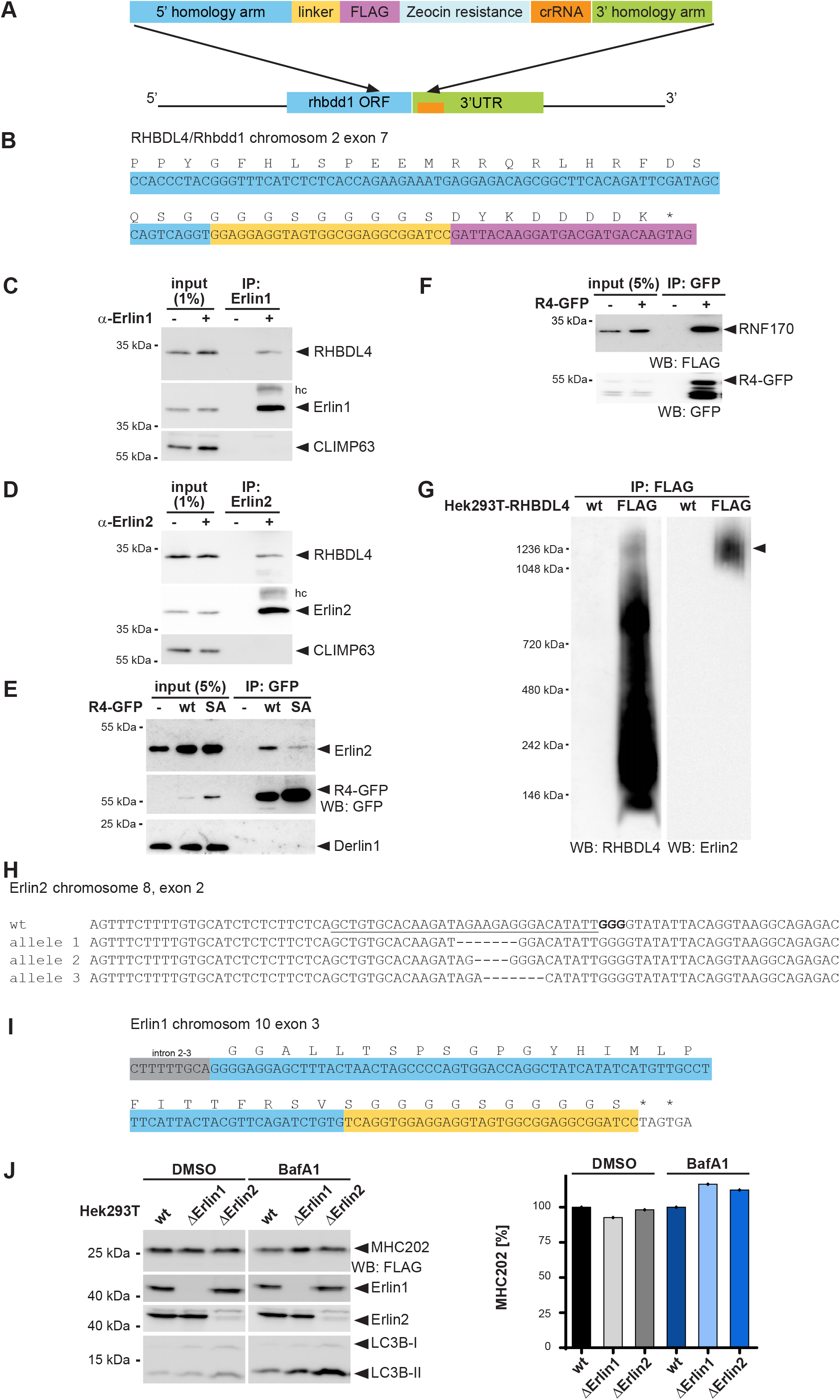
RHBDL4 interacts with Erlin1, Erlin2 and RNF170. **(A)** Outline of the applied tagging strategy of RHBDL4 (referred to by its gene name *Rhbdd1*) according to (Fueller et al., 2020). **(B)** Sanger sequencing of chromosomal DNA obtained from the Hek293T-R4-FLAG cell lines shows the insertion of the FLAG tag in the last coding exon. Colour code as in (A). **(C)** Microsomes from Hek293T cells were solubilized with 1% Triton X-100, and endogenous Erlin1 was isolated by immunoprecipitation (IP). Western blot (WB) identifies co-purification of endogenous RHBDL4. Hc, heavy chain. **(D)** Microsomes from Hek293T cells were solubilized with 1% Triton X-100, and endogenous Erlin2 was isolated by immunoprecipitation (IP). WB identifies co-purification of endogenous RHBDL4. Hc, heavy chain. **(E)** Hek293T cells were transfected with empty vector, RHBDL4-GFP or catalytic inactive RHBDL4-SA-GFP (SA). Following solubilization with Triton X-100, R4-GFP was isolated by immunoprecipitation (IP) using an anti-GFP antibody. Endogenous Erlin2 binds more efficiently to R4-GFP wild type (wt) than to its catalytic inactive SA mutant. Derlin1 was used as a negative control. **(F)** Co-immunoprecipitation experiment as in (C) using Hek293T cells transfected with RNF170-FLAG. **(G)** Endogenous RHBDL4 is part of an MDa-sized erlin complex. Hek293T wt cells or cells with chromosomally FLAG-tagged RHBDL4 (FLAG) were solubilized with 1% Triton X-100, immunoprecipitated for FLAG, eluted with FLAG peptides and analyzed by BN-PAGE. RHBDL4-FLAG formed several higher molecular weight complexes in addition to the 1.2 MDa complex containing Erlin2 (filled triangle). **(H)** Sanger sequencing of genomic DNA obtained from Hek293T Erlin2 knockout cells. Single-guide RNA-binding site is underlined; protospacer-associated motif is shown in bold. **(I)** Sanger sequencing of genomic DNA obtained from Hek293T Erlin1 knockout cells show the insertion of the Stop cassette in exon 3. Colour code as in (A). Erlin1/Erlin2 double knockout cells were generated accordingly using the Erlin2 knockout cells in (H). **(J)** Steady-state levels of MHC202 are reduced in Hek293T Erlin1 (ΔErlin1) and Erlin2 (ΔErlin2) knockout cells compared to wt Hek293T cells. Lysosomal inhibition by BafA1 increases stabilization of MHC202 caused by single knockout of Erlin1 and Erlin2 (n=1). Data information: For clarity, for panels C-G representative experiments of three independent replicates are shown.

**Figure 6 – figure supplement 1.**
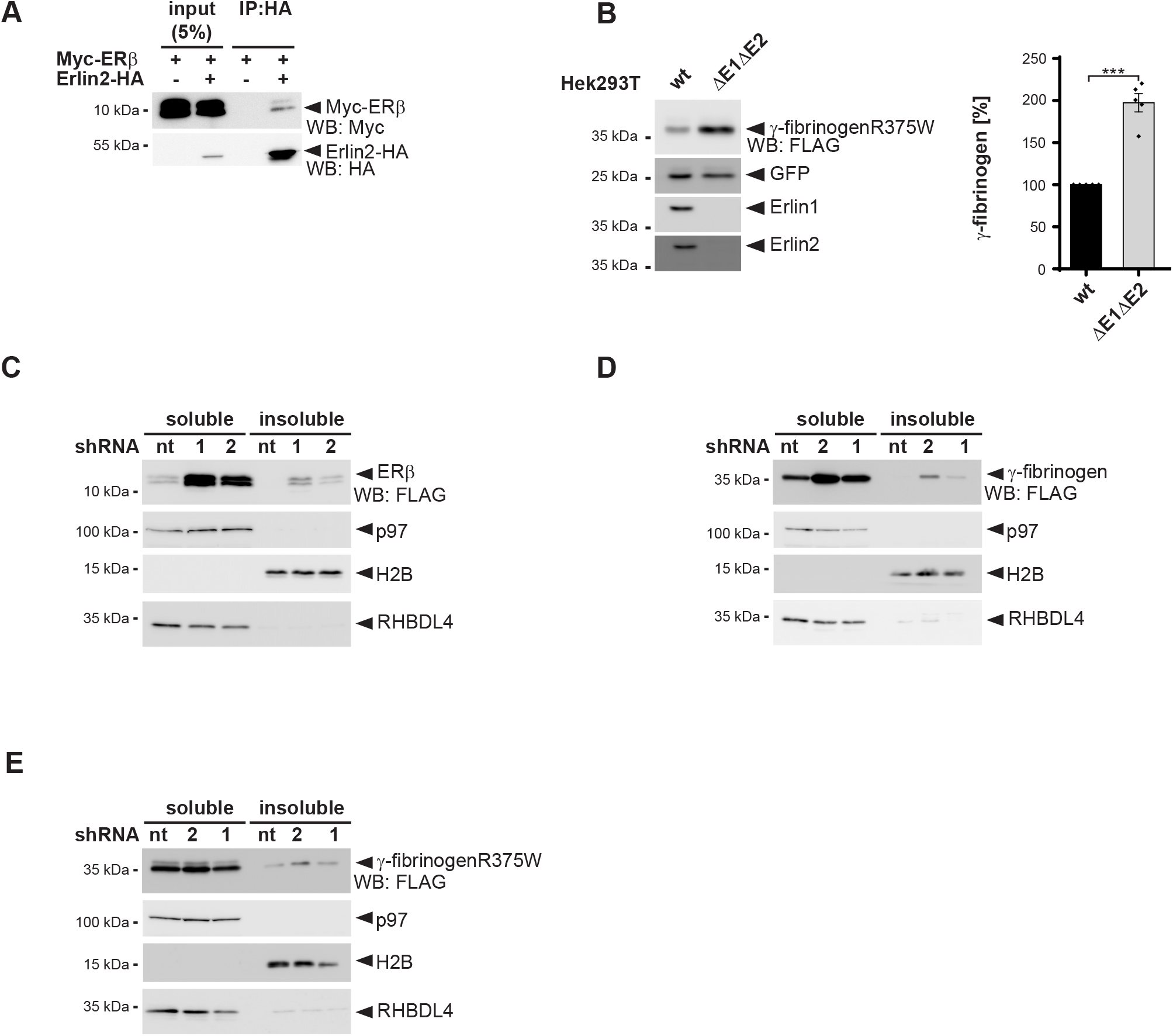
The RHBDL4-erlin complex interacts with aggregation-prone proteins. **(A)** Western blotting (WB) after immunoprecipitation (IP) of Erlin2-HA from Triton X-100 solubilized Hek293T cells confirms interaction with ERβ as has been shown previously (Vincenz-Donnelly et al., 2018). **(B)** Steady-state levels of γ-fibrinogen R375W mutant increase in Erlin1/Erlin2 Hek293T double knockout cells (ΔΔE1E2) in comparison to parental wild-type Hek293T cells (wt). GFP was used as a loading control. Right panel, WB quantification of γ-fibrinogen R375W (means ± SEM, n=5, ∗∗∗p≤ 0.001 (Student’s t-test)). **(C)** ERβ steady-state levels increase in Hek293T cells transfected with two independent shRNAs targeting RHBDL4 (R4-1 and R4-2) compared to non-targeting control (nt). Knockdown of RHBDL4 further increases the recovery of ERβ in the NP40 insoluble fraction. p97 was used as a loading control for the soluble fraction and H2B for the insoluble fraction. **(D)** γ-fibrinogen steady-state levels increase in Hek293T cells transfected with two independent shRNAs targeting RHBDL4 (R4-1 and R4-2) compared to cell treated with non-targeting control (nt) in Hek293T cells. Knockdown of RHBDL4 further increases the recovery of γ-fibrinogen in the NP40 insoluble fraction. p97 was used as a loading control for the soluble fraction and H2B for the insoluble fraction. **(E)** Assay as shown in (E) analysing the steady-state levels of the γ-fibrinogen R375W mutant. Knockdown of RHBDL4 further increases the recovery of γ-fibrinogen R375W in the NP40 insoluble fraction. p97 was used as a loading control for soluble fraction and H2B for insoluble fraction, respectively. Data information: For clarity, for panels A and C-F representative experiments of three independent replicates are shown. For panel B a representative experiment of five independent replicates is shown.

